# Virus-like transposons cross the species barrier and drive the evolution of genetic incompatibilities

**DOI:** 10.1101/2022.07.12.499685

**Authors:** Sonya A. Widen, Israel Campo Bes, Alevtina Koreshova, Daniel Krogull, Alejandro Burga

## Abstract

Horizontal gene transfer—the movement of genetic material between different species—has been reported across all major eukaryotic lineages, including vertebrates. However, the underlying mechanisms of transfer and their impact on genome evolution are still poorly understood. While studying the evolutionary origin of a selfish element in the nematode *C. briggsae*, we discovered that *Mavericks*, ancient viral-like transposons related to giant viruses and virophages, are one of the long-sought vectors of horizontal gene transfer. We found that *Mavericks* gained a novel herpesvirus-like fusogen in nematodes, leading to the widespread exchange of cargo genes between extremely divergent species, bypassing sexual and genetic barriers spanning hundreds of millions of years. Our results show how the union between viruses and transposons—nature’s melting pot—causes horizontal gene transfer and ultimately genetic incompatibilities in natural populations.

## Main Text

Genes, like early explorers who braved the seas, can also embark on extraordinary journeys. Horizontal gene transfer (HGT)—the non-sexual movement of genetic material between different species—is a well-characterized and common phenomenon in prokaryotes [1]. In contrast, transfer of genes between animal species is thought to be rare, as it requires a chain of unlikely events. First, DNA must find its way out of the donor species, come in close contact with the germline of a second species, and finally integrate itself in the genome of the new host. Nonetheless, from antiparasitic toxins in butterflies to antifreeze proteins in fish, a growing body of evidence indicates that HGT between animals is far more common than anticipated and could be an important force in eukaryotic evolution [2–11]. Yet one of the most fascinating aspects of HGT—the mechanism of transfer—remains a mystery. Viruses have been proposed to mediate HGT due to their known ability to take up sequences from their hosts and integrate into genomes [12–16]. However, evidence is sparse and mostly limited to highly specialized viral symbionts that have co-evolved in the genomes of parasitoid wasps for millions of years [12,17–20]. Here, by studying the evolutionary origin of a selfish toxin-antidote element (TA) in the nematode *Caenorhabditis briggsae*, we uncover *Mavericks*, ancient eukaryotic virus-like transposons, as widespread vectors of HGT between nematodes and show how HGT fueled the evolution of genetic incompatibilities in natural populations.

### A maternal-effect selfish TA in *C. briggsae*

TAs are made up of two linked genes, a toxin and its cognate antidote. In a maternal-effect TA, the toxin is expressed in the germline of mothers and loaded into all eggs prior to fertilization [21]. Only embryos that inherit at least one copy of the TA can express the antidote to counteract the effect of the toxin and survive [22]. In this way, TAs selectively decrease the fitness of non-carriers and rapidly spread in populations. We recently discovered that a genetic incompatibility between the *C. briggsae* reference strain AF16 (Ahmedabad, India) and the wild isolate HK104 (Okayama, Japan) stems from a maternal-effect TA [23–25]. This TA causes developmental delay in ∼20% of their F_2_ progeny, and all delayed progeny are homozygous non-carrier (AF16) individuals (Figure 1A). When grown at 25°C, *C. briggsae* typically takes two days to develop from embryos to sexually mature adults; however, non-carrier individuals are unable to counteract the effect of the toxin and take three days or longer to reach sexual maturity [23,24]. While the delay locus was previously mapped to HK104 Chr. III, the toxin-and antidote-encoding genes remain unknown.

**Figure 1.**
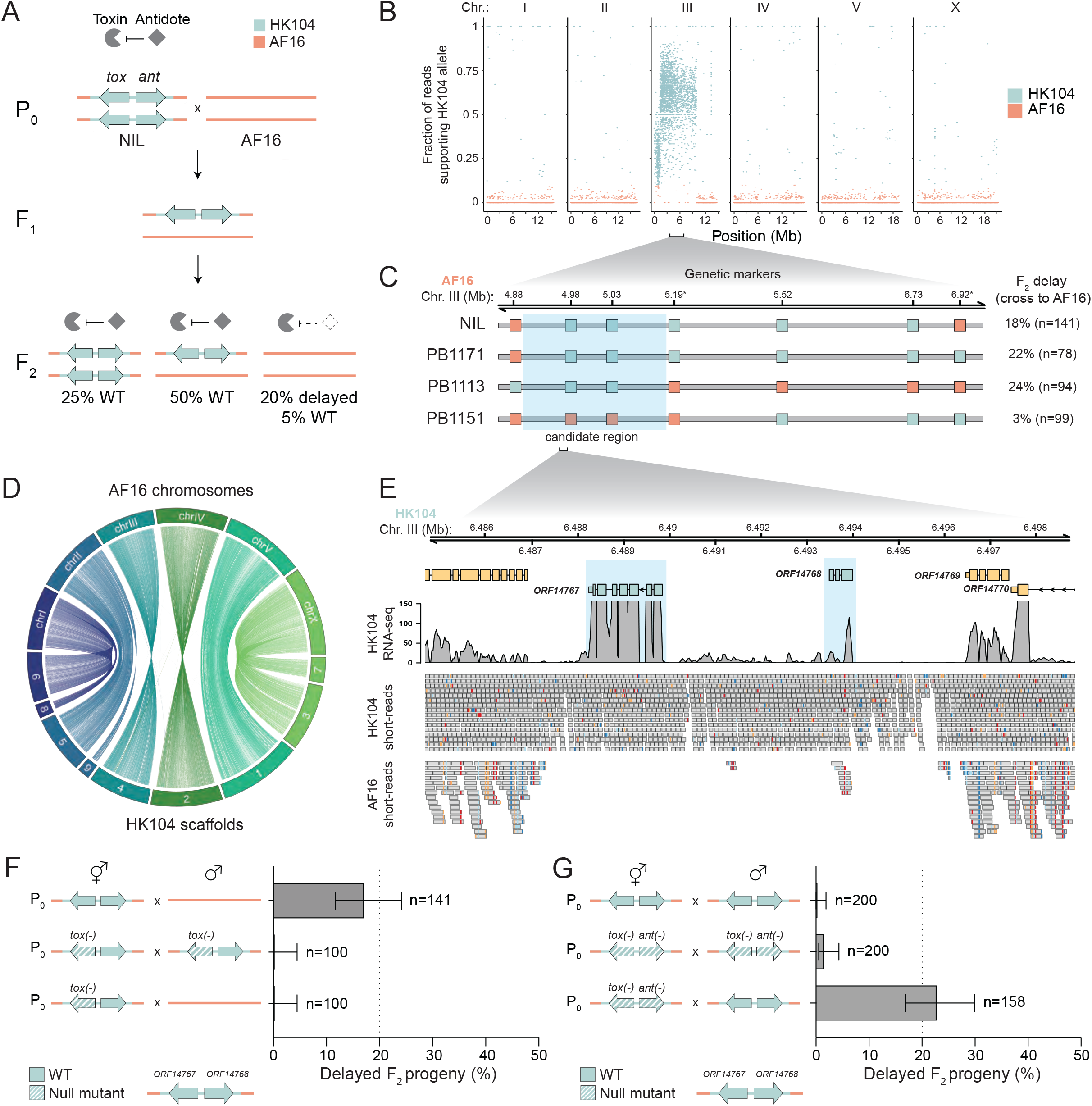
A maternal-effect selfish TA in *C. briggsae*. **(A)** Model illustrating the mechanism of action of the TA identified in HK104. In crosses between a hypothetical near-isogenic line (NIL) carrying the TA and the non-carrier strain (AF16), approximately 20% of the F_2_ progeny are developmentally delayed because AF16 homozygotes have not inherited the TA, and therefore cannot express the antidote to counteract the maternally-deposited toxin. The F_2_ delay phenotype is not fully penetrant and a small proportion of AF16 homozygotes (∼5% of F_2_ progeny) develop normally [23]. **(B)** Allele frequencies after 16 generations of maternal backcrosses to AF16, illustrating a large introgression on Chr. III. **(C)** Genotype of NIL1 and AF16 x HK104 recombinant interbred lines (RILs; PB1171, PB1113 and PB1151) at genetic markers across the Chr. III introgression to narrow the candidate region (given in AF16 coordinates). To test if each line still carries the HK104 TA, they were crossed to the non-carrier strain AF16 and their F_2_ progeny were analyzed for developmental delay (right). Asterisk (*) indicates genetic marker information that was gathered from Ross et al. [24]. **(D)** Synteny analysis between *C. briggsae* reference strain AF16 chromosomes and the HK104 *de novo* assembly scaffolds. The 20 HK104 scaffolds are numbered according to size (largest to smallest) and the 11 remaining much smaller scaffolds are not shown. **(E)** Alignment of short-reads and mixed embryo-stage RNA-seq to HK104 *de novo* assembly (y-axis is coverage). Highlighted in blue are *ORF14767* and *ORF14768*, two candidate genes present and expressed in HK104 but absent in AF16. **(F)** Genetic dissection of the toxin. Compared to the F_2_ developmental delay seen in crosses between the NIL and AF16 (top), a frameshift mutation in *ORF14767* in the NIL background is sufficient to completely rescue the delay phenotype in F_2_ progeny (bottom), indicating that *ORF14767* encodes the toxin. **(G)** Genetic dissection of the antidote. Introduction of an *ORF14768* frameshift mutation in the toxin mutant background does not itself cause developmental defects (middle). However, *ORF14767(-) ORF14768(-)* double mutants behave as a non-carrier strain and F_2_ delay phenotypes are fully restored in crosses with the wildtype NIL (bottom), indicating that *ORF14768* encodes the antidote. In all panels, HK104 is shown in cyan, AF16 is shown in salmon. Error bars indicate 95% confidence intervals calculated with the Agresti-Coull method. tox=toxin, ant=antidote.

To fine-map the underlying genes, we took advantage of the inherent gene-drive activity of TAs [23]. We backcrossed hybrid hermaphrodites to non-carrier AF16 males for 16 generations and sequenced the genome of the resulting population (Figure S1A; Methods). We expected the genetic background of the backcrossed population to be AF16 except for the HK104 Chr. III region that contains the TA. Consistent with this prediction, we found a single introgression in the left/central region of Chr. III, which spanned approximately 10 Mb (Chr. III: 0-10 Mb; Figure 1B). To narrow down the candidate region, we derived 50 independent near-isogenic lines (NILs) from the backcrossed population (Figure S1A) and genotyped them using established markers [26,27]. From these lines, we selected two NILs carrying large but partially overlapping introgressions, the exact coordinates of which we determined with Illumina short-read sequencing (overlap region: 4.94-6.75 Mb; Figure S1B). Using crosses we determined that the TA was located in the overlapping introgressed region (Figure S1C; Table S1). Repeated attempts to narrow the introgressions via backcrossing were unsuccessful, likely due to the low rate of recombination in the center of *C. briggsae* chromosomes [24]. To circumvent this, we took advantage of existing AF16 x HK104 recombinant interbred lines (RILs) and selected three RILs carrying different breakpoints across the candidate region [24]. By crossing each RIL to AF16 and testing for delay in the F_2_ population, we narrowed the candidate region to 151 kb (Chr. III: 4.94-5.09 Mb; Figure 1C). For all future experiments, we used NIL1 with the smallest introgression (hereafter simply referred to as NIL) and confirmed that it behaves similarly to HK104 in both maternal and paternal backcrosses to AF16 (Figure S1D).

To identify candidate genes, we *de novo* assembled the HK104 genome using a combination of Oxford Nanopore long-reads and Illumina short-reads. The final assembly consisted of 18 scaffolds, with an N50 of 14,833,495 bp (Figure 1D). We then sequenced the HK104 transcriptome and created comprehensive gene annotations using both transcriptome-guided and *de novo* gene calling. Our final annotation included 26,658 genes (27,569 transcripts), and analysis of conserved genes showed very high completeness (96.7% of conserved nematode genes were identified using BUSCO). Among the 39 genes in the candidate region, we filtered for genes in HK104 that were either (1) absent, mutated or highly divergent in AF16, or (2) lowly expressed in AF16, and (3) in tight genetic linkage (usually close neighbors). Only one pair of genes satisfied these criteria: *ORF14767* and *ORF14768* (Figure 1E; Table S2).

Due to its high expression level, we hypothesized that *ORF14767* is the toxin-encoding gene. To test this, we first used CRISPR/Cas9 homology-directed repair in the NIL background to introduce a premature stop codon in the second exon of *ORF14767*, leading to early truncation of the predicted protein (p.S77X; Figure S2). During the generation of these worms, *ORF14767* was duplicated but the second locus codes for an identical protein (Figure S2). The resulting *ORF14767(-)* worms did not show any abnormal phenotypes (Figure 1F). To test if the *ORF14767* mutation disrupted toxicity of the TA, we crossed *ORF14767(-)* NIL hermaphrodites to AF16 males. We observed no delayed individuals among the F_2_ progeny (n=0/100; Figure 1F), compared to 17.8% in the *ORF14767* wildtype cross (Figure 1C), indicating that *ORF14767* is the HK104 toxin. The toxin encodes an uncharacterized protein (325 amino acids long) with a predicted serine-protease domain (Figure S3).

Finally, we determined that *ORF14768* encodes the antidote. To show this, we first mutated *ORF14768* in the background of the toxin mutant using CRISPR/Cas9 (p.E87GfsX2; Figure S4); the resulting worms showed no abnormal phenotypes (Figure 1G). We then crossed *ORF14767(-) ORF14768(-)* double mutant hermaphrodites to the parental NIL males. If these two genes respectively encode the toxin and the antidote, we would expect the double mutant to be functionally equivalent to a non-carrier AF16 allele, which lacks both. As expected, F_2_ developmental delay was fully restored to wildtype levels (n=36/158, 22.8%; Figure 1G), and essentially all delayed individuals were homozygous double mutants (n=34/36; Table S1). We further recapitulated these data with an independently-derived *ORF14768(-)* in the toxin mutant background (p.K65X; Figure S4) (n=19/80, 23.8%; Table S1) confirming that *ORF14768* is the antidote-encoding gene. The antidote encodes an uncharacterized small protein—only 153 amino acids long—that has no predicted protein domains (Figure S3).

### *msft-1/tlpr-1* is a novel mobile selfish TA element

To study the evolutionary origins of the TA, we focused our attention on the surrounding sequences. Although our annotation pipeline did not identify any clear open reading frame in the ∼3.7 kb region between the toxin and the antidote, we detected low levels of transcription (Figure 1E). By leveraging *de novo* gene predictions and evolutionary conservation, we identified a gene with high homology to transposases belonging to the MULE (<u>Mu</u>tator-like <u>e</u>lements) family of transposable elements (Figure 2A; Figure S5). MULEs are “cut-and-paste” DNA transposons that are known for their propensity to insert into genic regions and acquire host genes [28–30]. Remarkably, the toxin, the MULE transposase, and the antidote were flanked by terminal inverted repeats (TIRs)—870 bp long and 96.7% identical—strongly suggesting not only a functional association between the TA and the MULE, but also a recent transposition event. In light of this, we named the toxin *msft-1* (for *<u>M</u>aternal <u>S</u>erine-protease toxin <u>F</u>eaturing <u>T</u>ransposon*), and the antidote *tlpr-1* (for *<u>T</u>heriac against <u>L</u>arval delay and <u>PR</u>otease toxicity*). To test whether the *msft-1/tlpr-1* TA is indeed mobile in nature, we assembled the genomes of three additional *C. briggsae* wild isolates using Nanopore long-reads (Figure S6). Analysis of these three and two additional genomes [31] revealed that the insertion site of *msft-1/tlpr-1* varied between natural isolates. *msft-1/tlpr-1* was located on Chr. III in wild isolates from China and Iceland but it was found on Chr. V in isolates from Canada, Saint Lucia, and Taiwan (Figure 2B,C). The Chr. V TA was pseudogenizeded but largely complete. It lacked the transposase and antidote sequences but it contained most of the toxin and fragments of both flanking MULE TIRs. Thus, standing variation in wild *C. briggsae* populations supports the view that the *msft-1/tlpr-1* TA was capable of excision and integration into new genomic loci in recent evolutionary history.

**Figure 2.**
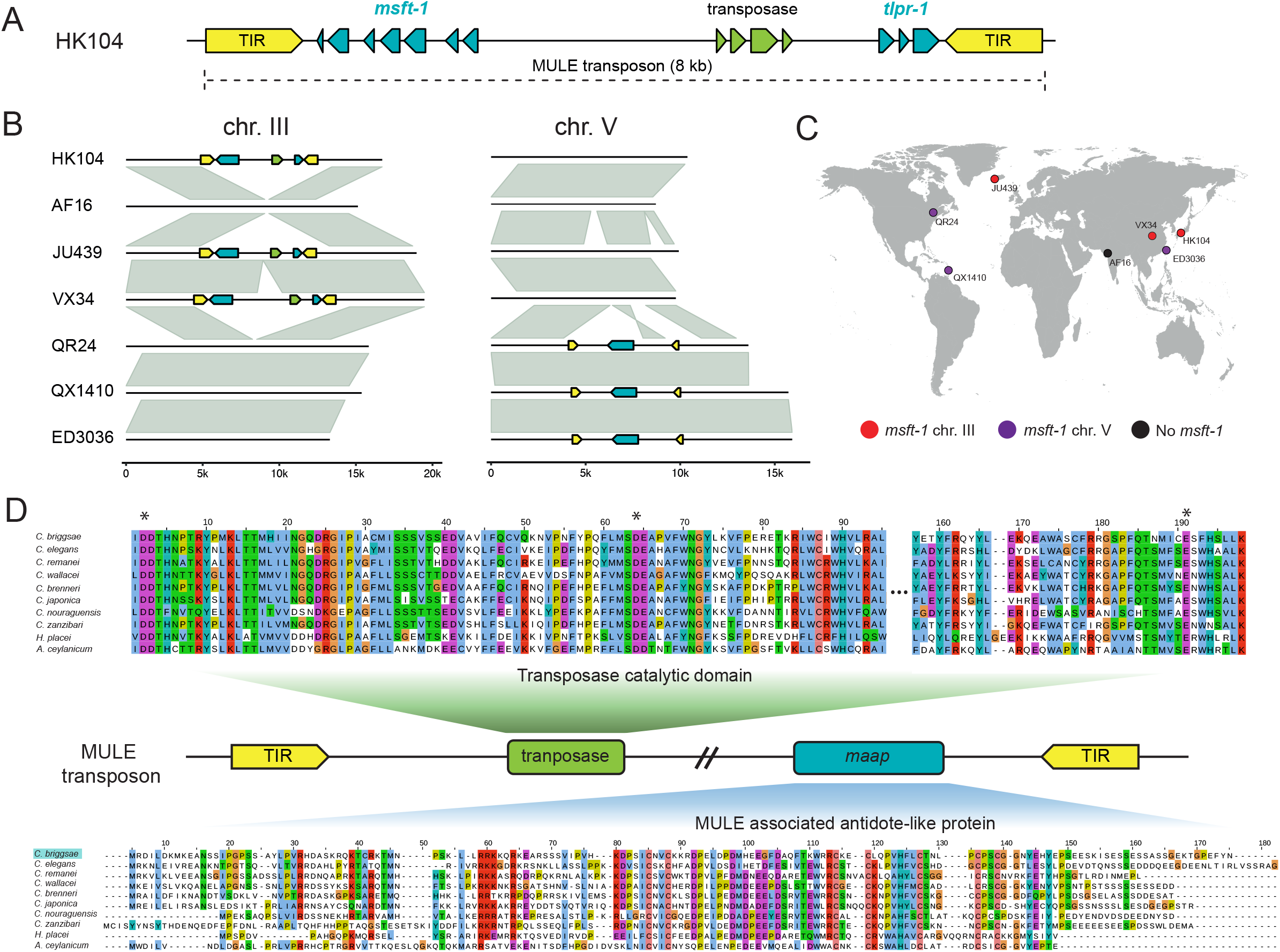
*msft-1/tlpr-1* is a mobile selfish TA. **(A)** The *msft-1/tlpr-1* TA is contained within a MULE transposon. A conserved MULE transposase is found in between the toxin and the antidote and all three genes are flanked by TIRs (terminal inverted repeats). **(B)** Natural genetic variation in *C. briggsae* populations supports the mobilization of the *msft-1/tlpr-1* TA in wild populations. Alignment of Chr. III and Chr. V regions that carry a copy of the TA. **(C)** Sampling distribution of *C. briggsae* wild isolates. Color code shows the TA localization. **(D)** Conservation of nematode MULE transposons. MULE transposase (green) and MAAP (MULE-associated antidote-like protein) (cyan) are flanked by TIRs (yellow). Alignment of representative MULE proteins across nematodes. Asterisks (*) show catalytic residues in the transposase.

### The *tlpr-1* antidote was co-opted from a MULE transposon

As well as their conserved transposase, MULEs typically contain additional ORFs [32]. For instance, the highly studied *Mu* transposon in Maize is made up of two genes: the *mudrA* transposase and a second small gene, *mudrB*, which has been proposed to mediate transposon integration [33]. Unlike their counterparts in plants, MULEs are poorly characterized in nematodes. To fill this gap, we first predicted the structure of the *C. briggsae* MULE transposase using AlphaFold2 [34] and identified the residues that make up its characteristic DDE motif catalytic triad based on structural homology to IS*Cth4*, a prokaryotic transposase of the IS256 family that is related to eukaryotic MULEs (Figure S7) [35]. These three residues (Asp21, Asp83 and Glu198) were perfectly conserved in MULE transposases from 49 different nematodes species, including HK104, suggesting that the transposase linked to *msft-1* is functional (Figure 2D). We then annotated orthologs of *tlpr-1* across a wide range of divergent nematodes, including distantly related parasitic species such as *Haemonchus placei* and *Ancylostoma ceylanicum* (Figure 2D). Surprisingly, *tlpr-1* homologs were found immediately downstream of the transposase catalytic domain in ∼40% (67/161) of annotated MULE transposons, suggesting a functional association between the two genes. We thus named the new protein family MAAP (for <u>M</u>ule-<u>A</u>ssociated <u>A</u>ntidote-like <u>P</u>rotein). Except for the *msft-1/tlpr-1* locus, MULE transposons were not associated with *msft-1* in any of the nematode species examined. Therefore, our results strongly suggest that the *msft-1/tlpr-1* TA was not captured by a MULE, but that the MULE first captured *msft-1* and only then the MULE-associated protein was co-opted into its cognate antidote, becoming *tlpr-1*.

### The MSFT-1 toxin belongs to a novel family of serine proteases

To better understand the mechanism underlying MSFT-1 toxicity, we predicted its tertiary structure using AlphaFold2 [34]. The resulting model had a high accuracy (Figure S8) and revealed two distinct domains that are connected by a flexible linker (Figure 3A). The N-terminal domain [AA1:83] displays a ubiquitin-like fold, whereas the C-terminal one [AA:106-325] matches the serine protease domain predicted by InterPro (Figure S3A). An independent model generated using RoseTTAfold resulted in a similar structure (RMSD=1.3Å and TM-score=0.85 for the protease domain; Figure S8) We then used the AlphaFold2 MSFT-1 model as a template to search for proteins with a similar structure in the Protein Data Bank (PDB) using DALI [36]. MSFT-1 was similar to the HtrA (<u>H</u>igh-temperature requirement <u>A</u>) family of serine proteases (Figure 3B) [37–39]. HtrA proteases are widespread in bacteria and eukaryotes and they typically contain one C-terminal PDZ domain, which stabilizes interactions with their substrates and modulates the activity of the protease domain [40]. In bacteria, HtrA proteases—also known as Deg—mediate protein quality control and can also act as virulence factors [39,41,42]. Despite sharing very low sequence similarity and lacking a PDZ domain, DELTA-BLAST and HHbit—two remote homology detection algorithms— identified MSFT-1 as homologous to bacterial Deg proteases. We then aligned the protease domain of MSFT-1 to *Pseudomonas aeruginosa* AlgW, a representative HtrA protein (PDB: 7CO5), and identified the residues that make up its catalytic triad based on structural homology: His148, Asp184, and Ser256 (RMSD=1.9Å; TM-score=0.71; Figure 3C; Figure S9). To test whether the protease activity of MSFT-1 is necessary for its toxicity, we mutated Ser256 to alanine in the endogenous *msft-1* locus using CRISPR/Cas9. This mutation abrogates the activity of serine proteases because it renders them incapable of performing the initial nucleophilic attack [43]. We then crossed *msft-1* Ser256Ala NIL hermaphrodites to AF16 males and inspected their F_2_ progeny. In contrast to crosses involving the wildtype *msft-1/tlpr-1* TA, we observed no developmental delay among F_2_ individuals (Figure 3D), and all F_2_ homozygous non-carrier (AF16) worms were phenotypically wildtype. These results were further confirmed in crosses with an independently derived line carrying the same point mutation (0.55% F_2_ delay, n=1/180). Thus, our results indicate that the proteolytic activity of MSFT-1 is essential for its toxicity *in vivo*.

**Figure 3.**
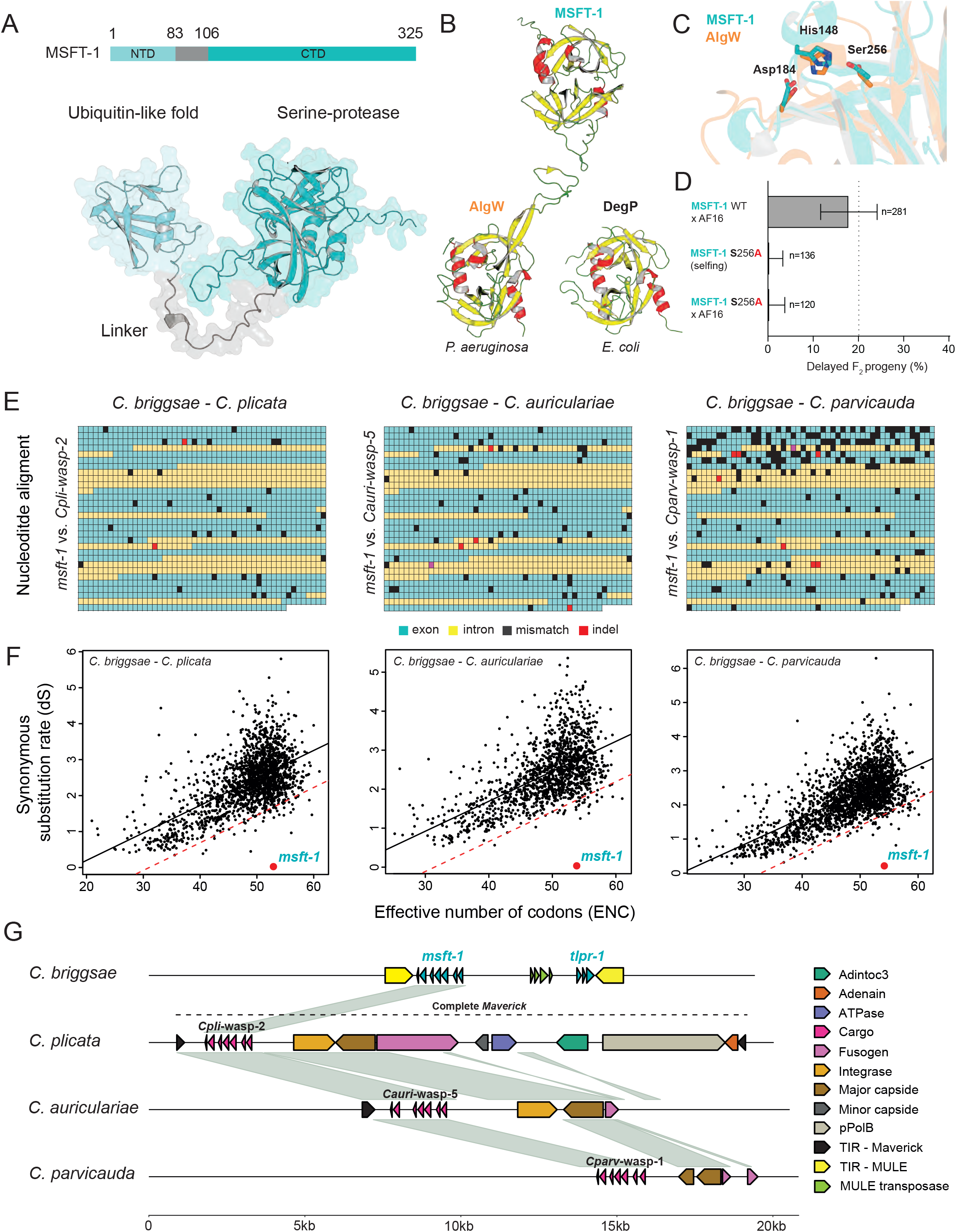
Horizontal gene transfer of *msft-1* and related proteases. **(A)** Predicted domains and AlphaFold2 model of MSFT-1. **(B)** Comparison of the serine-protease domain of MSFT-1 and two representative HtrA proteases, AlgW from *Pseudomonas aeruginosa* (PDB: 7CO5) and DegP from *Escherichia coli* (PDB: 6JJK). α-chains in red, β-sheets in yellow, linkers in green. **(C)** Structural alignment of the protease domains of MSFT-1 (cyan) and AlgW (orange). Focus is on the three key catalytic residues. **(D)** Mutation of MSFT-1 Ser256 to Ala abolishes its toxicity in crosses. Error bars indicate 95% confidence intervals calculated with the Agresti-Coull method. **(E)** Nucleotide alignment of *C. briggsae msft-1* to its closest orthologs: *C. plicata wasp-2, C. auriculariae wasp-5*, and *C. parvicauda wasp-1*. Alignments include intronic sequences. Mismatches, deletion and insertion are highlighted. **(F)** Vertical and Horizontal Inheritance Consistence Analysis (VHICA) for each species pair. Number of single copy orthologs analyzed for each species pair: *C. briggsae* HK104 and *C. auriculariae* (1370); C. briggsae HK104 and C. *plicata* (1775); and *C. briggsae* HK104 and *C. parvicauda* (2172). *msft-1*/*wasp* gene pairs are highlighted in red circles. **(G)** Alignment of the *msft-1/tlpr-1* TA and its closest orthologous *wasp* proteases in basal *Caenorhabditis* species. Highlighted are genes and TIRs of MULE and *Maverick* transposable elements. The *Maverick*s of *C. auriculariae* and *C. parvicauda* are pseudogenized.

### Horizontal gene transfer of *msft-1* and related proteases

To further investigate the evolutionary history of MSFT-1, we retrieved homologous sequences from 107 nematode genomes, curated gene annotations, and built a comprehensive phylogenetic tree (Figure S10; Data S1). In total, we identified 235 homologous proteins distributed across 53 species and 11 genera. To facilitate their study, we named this novel family of proteases WASP (for *<u>W</u>orm Htr<u>A</u>-like <u>S</u>erine <u>P</u>rotease;* see Data S1). Our analysis revealed that WASP proteases are very common in nematodes but not universal. For instance, *C. briggsae* has two WASP proteases, CBG29787 and CBG26403, whereas *C. elegans* lacks WASP orthologs altogether. While inspecting the tree branch containing the toxin in detail, we made an intriguing observation. MSFT-1 was almost identical to *Cpli*-WASP-2, a protease found in *C. plicata*, a rare *Caenorhabditis* species associated with carrion beetles [44]. Their sequences were 95% identical at the amino acid level (Figure S11). This is surprising because *C. plicata* is one of the most basal species of the *Caenorhabditis* genus (Figure S12) [45]. More strikingly, their coding and intronic sequences were 97.5% identical at the nucleotide level, even though the genomes of *C. briggsae* and *C. plicata* are at least as divergent as those of humans and zebrafish (Figure 3E; Figure S13) [46]. In addition, we found high nucleotide conservation between MFST-1 and WASP proteases found in *C. parvicauda* (*Cparv*-WASP-1) and *C. auriculariae* (*Cauri-*WASP-5), two basal *Caenorhabditis* species, and *C. inopinata* (*Cino-*WASP-2) (Figure 3E), the sister species of *C. elegans* (Figure S14) [45,47,48]. These observations raised a critical question— how can MSFT-1 be almost identical to orthologs in species that last shared a common ancestor well over 100 million years ago?

Introgressive hybridization—the exchange of genetic material between different species via hybridization and backcrossing—can result in high levels of identity between orthologs [49]. However, this phenomenon is unlikely to explain the extreme sequence identity of *msft-1* for two main reasons. First, with the exception of *C. nigoni*, there is complete reproductive isolation between *C. briggsae* and all other *Caenorhabditis* species [50]. Second, such high levels of sequence identity could only be the result of a recent hybridization event. If the introgression were recent, we would expect to find extensive homology in the vicinity of the *msft-1* locus due to linkage disequilibrium [51]. However, that was not the case; only the protease had a high sequence identity, while the neighboring genomic regions were not homologous (Figure S15). Because such an isolated and extreme case of sequence similarity was unlikely to arise by hybridization, we hypothesized that *msft-1* has been horizontally transferred.

To test whether *msft-1* has been horizontally transferred, we studied its phylogeny in more detail. A landmark of horizontal gene transfer (HGT) is the presence of significant discrepancies between species and gene trees [52]. In addition to the high sequence similarity between *C. briggsae msft-1* and orthologs found in *C. plicata, C. parvicauda, C. auriculariae* and *C. inopinata*, we found that basal *Caenorhabditis* species and those belonging to the *Elegans* and *Japonica* groups were frequently intermixed in the protease tree (Figure S10). Furthermore, *Caenorhabditis* species no longer formed a monophyletic group (Figure S10). For example, *C. briggsae* WASP-1 was most closely related to *Pristionchus pacificus* WASP-1, *C. plicata* WASP-1 to *Mesorhabditis belari* WASP-12, and *C. virilis* WASP-1 to *Mesorhabditis belari* WASP-8 (Figure S10).

Because gene loss and duplication can lead to topological discrepancies between gene and species trees [53], we also tested the HGT hypothesis using a method that is independent from phylogenetic reconstruction—Vertical and Horizontal Inheritance Consistence Analysis (VHICA) [54]. VHICA is a statistical framework that compares genome-wide rates of neutral evolution in coding genes and identifies horizontally transferred genes based on their unusually low rate of synonymous substitutions (dS). VHICA accounts for variation in the number of synonymous sites under purifying selection by using the effective number of codons (ENC) as a proxy, to distinguish between codon selection and HGT. To implement VHICA, we first defined a set of high confidence single copy orthologs between *C. briggsae* HK104 and the three species carrying proteases most similar to *msft-1*—*C. plicata, C. auriculariae*, and *C. parvicauda*—and estimated dS and ENC for each of them. As expected, we observed an overall positive correlation between dS and ENC on a genome-wide scale (Figure 3F). We then identified genes that significantly deviated from this trend; that is, they displayed both a low dS and high ENC (low codon bias), a hallmark of HGT. Following these criteria, *msft-1* was the most extreme outlier genome-wide—strongly supporting a HGT event (Figure 3F).

In summary, 1) their extreme sequence similarity in highly divergent lineages (including introns), 2) the incongruence between species and gene trees, 3) their uneven distribution across nematode species, and 4) their unusual pattern of molecular evolution compared to vertically inherited genes, led us to conclude that *msft-1* and many other *wasp* proteases were horizontally transferred in nematodes. Furthermore, our results suggest that HGT events involving these proteases are common in nature and can even take place between species that diverged hundreds of millions of years ago.

### *Mavericks* transferred the precursor of *msft-1* into *C. briggsae*

Having established that *msft-1* is horizontally transferred in nematodes, we wondered by what mechanism the transfers were accomplished. One key finding that has emerged from various studies is that HGT usually involves transposable elements [7,8,16]. Since *msft-1* is embedded within a MULE in *C. briggsae*, we wondered whether this DNA transposon was responsible for the horizontal transfer of *msft-1*. To test this idea, we reconstructed the evolutionary history of nematode MULEs and searched for phylogenetic incongruencies that could indicate horizontal transfer. MULEs are quite challenging to annotate because they are highly divergent, oftentimes pseudogenized, and their transposase is normally silenced or expressed at very low levels. Thus, we focused our analysis on the catalytic domain of the transposase, which is by far the most conserved domain of the enzyme (Figure 2D). Our analysis revealed that, unlike WASP proteases, *Caenorhabditis* MULE transposases form an almost perfect monophyletic group (Figure S16). Moreover, the *C. briggsae* MULE transposase was most similar to the one found in its sister species, *C. nigoni* (Figure S16). Lastly, we asked whether the *C. briggsae* MULE transposase or its TIRs were highly conserved in any other species. Unlike *msft-1*, we found no evidence for high sequence conservation of the MULE in any other viral, prokaryotic or eukaryotic genome publicly available to date. Taking all of these results into consideration, we concluded that MULE transposons are unlikely to be involved in HGT of WASP proteases.

We next thought to inspect in closer detail the genomic neighborhood of the *Cpli-wasp-2* protease, the closest homolog of *msft-1*. Although the protease was found in a very short scaffold, a *blastx* search revealed that the region at the closest edge of the scaffold and adjacent to the protease shared homology to viral integrases (Figure S17). Intrigued by this result and to overcome the limitations of the highly fragmented *C. plicata* short-read assembly (N50: 58,285 bp), we re-sequenced *C. plicata* using Nanopore long-reads and *de novo* assembled its genome. The resulting assembly consisted of 161 scaffolds and had N50 of 4,459,858 bp. Most genes were complete in this assembly despite the high levels of heterozygosity in the strain that hindered its assembly (BUSCO score 84.49% complete and 7.92% partial genes; Figure S17). The new long-read-based assembly revealed that *Cpli-wasp-2* is found within a complete *Maverick* transposon. In addition, both *Cauri-wasp-5* and *Cparv-wasp-1* are found within pseudogenized *Mavericks. Mavericks*—also known as polintons—are a class of ancient eukaryotic mobile elements that share features of both transposons and viruses [55,56]. Like transposons, *Mavericks* are flanked by TIRs. But like viruses, they code for a large number of proteins, including a type-B DNA polymerase, a retroviral-like integrase, a FstK-like ATPase, an adenovirus-like protease, as well as major and minor capsid proteins [57,58]. We found all of these intronless genes in the *Maverick* carrying *Cpli-wasp-2* (Figure 3G). The genetic link between *Mavericks* and giant viruses, virophages, and adenoviruses has led to the hypothesis that *Mavericks* could, under some conditions, form virions [58]. In support of this view, an AlphaFold2 model of the *C. plicata Maverick* major capsid protein (MCP) revealed extensive structural similarity to the double jelly-roll capsids of numerous dsDNA viruses, including the virophage *Sputnik*, as well as the giant viruses *Faustovirus* and *Paramecium bursaria* chlorella virus (PBCV-1) (top Z-score=24.4; Figure 4A-B). Viral MCP trimers—also known as hexons—are the main building block of capsids. The N-terminal tail of MCPs contribute to the stability of the homotrimer by wrapping around their nearest counterclockwise neighbouring subunit [59]. An AlphaFold2 model of the *Maverick* MCP trimer revealed that their N-terminal tail also shows this conserved structural feature (Figure 4A-B; Figure S18), suggesting that *Mavericks* can form virions.

**Figure 4.**
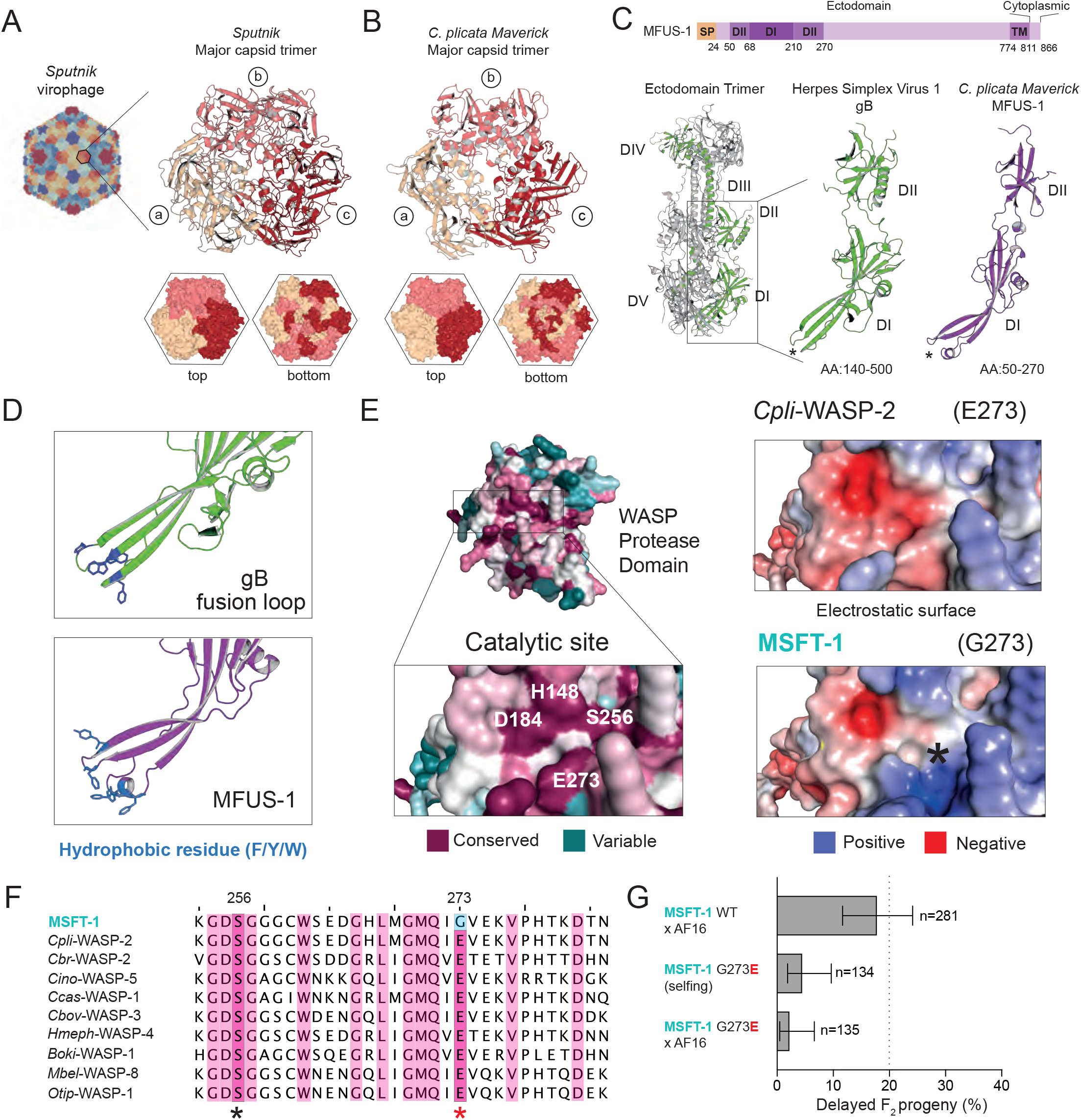
*Mavericks* transferred the precursor of *msft-1* into *C. briggsae*. **(A)** Structure of the *Sputnik* virophage capsid and major capsid protein trimer (hexon) (PDB: 3J26). **(B)** Predicted AlphaFold2 structure of *C. plicata Maverick* major capsid protein trimer. **(C)** Domain architecture of MFUS-1. Signal peptide (SP), transmembrane domain (TM) (top). Structure of the glycoprotein B ectodomain from HSV-1 (PDB: 2GUM). For simplicity, only one subunit of the homotrimer is shown in green. Key domains (DI-V) are highlighted (bottom left and middle). AlphaFold2 MFUS-1 model (bottom right). **(D)** Zoom in into the HSV-1 fusion loops in Domain I of glycoprotein B (top) and the homologous loops in the predicted MFUS-1 structure (bottom). Highlighted in blue are hydrophobic residues in the loops that are thought to interact with target membranes. **(E)** AlphaFold2 model of the serine-protease domain of *Cpli*-WASP-2. Colors represent the evolutionary conservation of each residue. Highlighted in white are the three catalytic residues and the highly conserved E273 (left). Electrostatic surface potential calculated for AlphaFold2 models of *Cpli*-WASP-2 (top right) and MSFT-1 (bottom right). The E273G substitution found in MSFT-1 alters the negative charge of the region next to the catalytic site (*). **(F)** Partial protein alignment of the representative MSFT-1 orthologs used to generate the surface plot in (E). Highlighted are residues S256 (catalytic serine) and E273 as shown in E. Highly conserved residues in magenta. **(G)** Reversal of MSFT-1 G273 near the catalytic site to the ancestral glutamic acid residue (G273E) abolishes its toxicity in crosses. Error bars indicate 95% confidence intervals calculated with the Agresti-Coull method.

Strikingly, the *C. plicata Maverick* additionally codes for a putative viral-like fusogen, which we named *mfus-1* (for *<u>M</u>averick <u>FUS</u>ogen*). Fusogens are proteins used by enveloped viruses to facilitate virus–host membrane fusion. *C. plicata* MFUS-1 is 866 amino acids long and, like most known viral fusogens, it has a predicted N-terminal signal peptide (AA:1-24) and a single transmembrane domain (AA:775-811) (Figure 4C). We used AlphaFold2 to predict the structure of MFUS-1 and focused our analysis on a portion of the ectodomain (AA:50-270) that showed the highest accuracy (Figure 4C; Figure S19). A structural homology-based search revealed that MFUS-1 was most similar to *Herpes simplex virus 1* (HSV-1) glycoprotein B (gB) (top Z-score=11.1; RMSD=3.88Å; TM-align 0.52; Figure 4C), and also showed extensive similarity to the glycoprotein B of human cytomegalovirus (HCMV) and Epstein-Barr virus (EBV), which are class III viral fusogens. Our approach identified the highly conserved domain I of class III proteins (DI), which forms an extended β-sheet structure and contains two loops that form the bipartite fusion peptide (Figure 4C). Like its viral counterparts, MFUS-1 has many hydrophobic residues in the putative loop regions of DI that could interact with target membranes (Figure 4D). Additionally, we found strong support for DII (Figure 4C) and DIII, a large α-helix that forms a trimeric coiled coil in viruses (Figure S19).

### Evolutionary path from protease to toxin

Since *Cpli-wasp-2* is embedded within a *Maverick* that could form enveloped virions, whereas MULE-associated *msft-1* can only directly mobilize within the *C. briggsae* genome, we reasoned that *Cpli-wasp-2* represents an ancestral state from which the toxic activity of *msft-1* evolved following a horizontal transfer event. To explore this scenario, we analyzed in detail the coding differences between these two genes. Of 16 non-synonymous substitutions between *Cpli-wasp-2* and *msft-1*, only one of them affects a highly conserved residue (E273; Figure 4E-F; Figure S11). This glutamic acid is found in very close proximity to the catalytic site of WASP proteases and is likely part of the substrate binding site (Figure 4E). In MSFT-1, however, this residue was replaced by a non-charged glycine (E273G; Figure 4E-F). To test whether this specific substitution is necessary for MSFT-1 toxicity, we first used CRISPR/Cas9 to revert G273 to the inferred ancestral state (glutamic acid) in the endogenous *msft-1* locus and then crossed these mutants to the non-carrier strain (AF16). We observed almost no delayed individuals among their F_2_ progeny (2.2%, n=3/135) indicating that this recent substitution is necessary for MSFT-1 toxicity (Figure 4G). We confirmed these results in crosses with an independently derived line carrying the same point mutation (5.6% delayed F_2_ progeny, n=9/159).

Overall, we conclude that *C. briggsae msft-1* evolved from a *Maverick*-associated protease that was horizontally transferred to *C. briggsae*, likely from a basal *Caenorhabditis* species where closely related homologs are found. *Mavericks* carry the genetic toolkit necessary for making enveloped virions, suggesting that viral-like particles could directly mediate HGT without the need of a vector, such as another virus or pathogenic microorganism.

### Widespread horizontal transfer of *Maverick* cargo across genera

MSFT-1 and its close homologs are not the only WASP proteases that are horizontally transferred; in fact, their patchy phylogeny suggests this is a common occurrence in the wild (Figure 5A; Figure S10). Prompted by our discovery in *C. plicata*, we next asked whether WASP proteases are generally associated with *Mavericks*. In 47 of 235 cases (20%), we identified at least one core component of *Mavericks* in close proximity (±6 kb) to a WASP protease (Figure 5A). We think this number is an underestimate, as the genome assemblies of most of these species are low-quality and the proteases are often found at the edge of scaffolds. WASP proteases were associated with *Mavericks* not only in *Caenorhabditis* species but also in distantly related nematodes such as *Mesorhabditis belari, Oscheius tipulae*, and even *Halicephalobus mephisto*, which was discovered inhabiting a South African mine 1.3 km below the Earth’s surface [60,61].

**Figure 5.**
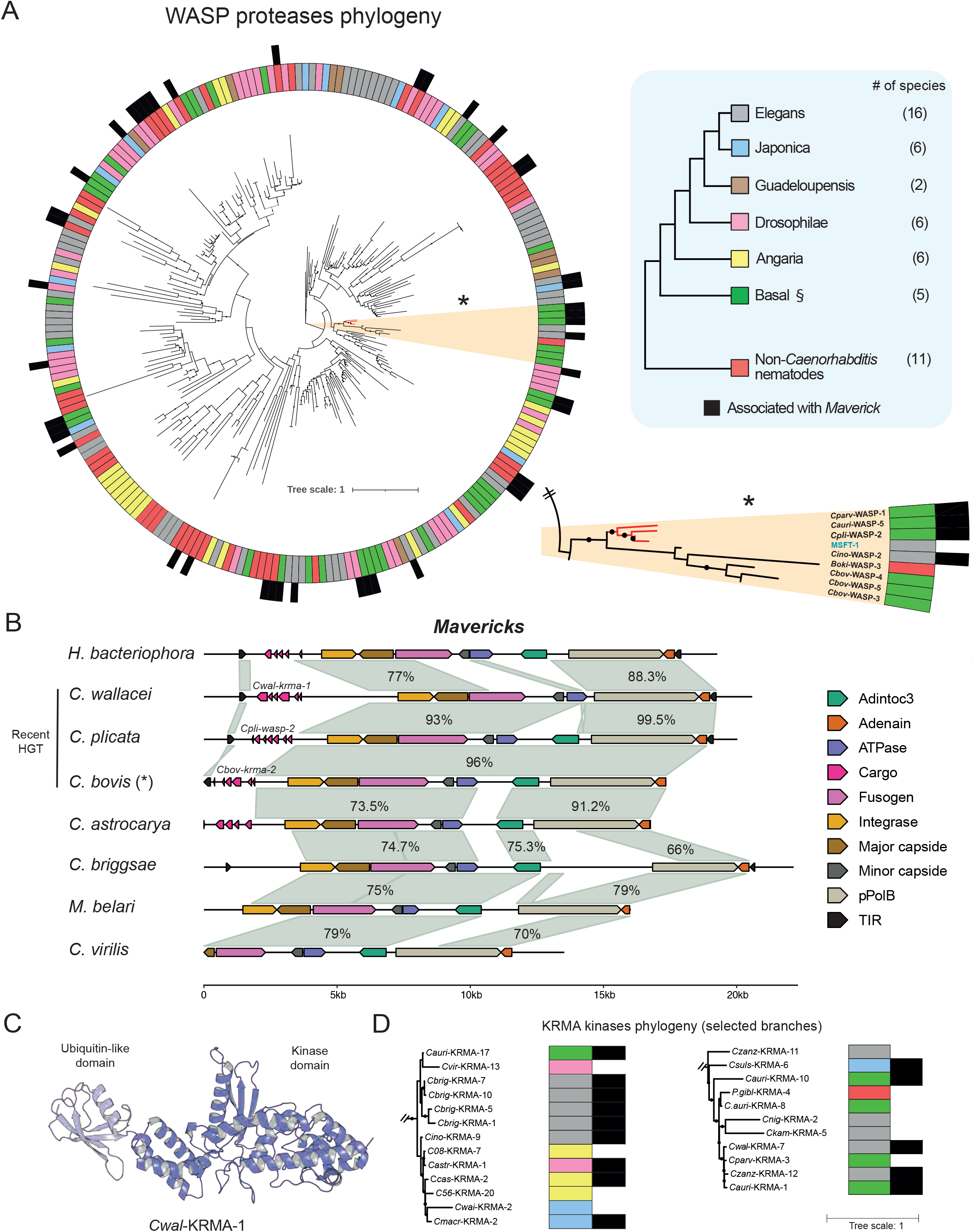
Widespread horizontal transfer of *Maverick* cargo across genera. **(A)** Protein phylogenetic tree of 235 WASP proteases found across 52 nematode species. The *Caenorhabditis* group to which each ortholog belongs is represented by the colour of the inner ring. These groups are monophyletic with the exception of the basal *Caenorhabditis* group (§) and the group containing non-*Caenorhabditis* species. Black outer ring represents those WASP proteases that are associated with *Mavericks* in the genome. Black dots denote branches with a bootstrap value >85. Zoom-in of the branch containing the MSFT-1 toxin. The closest orthologs in other species are coloured red. **(B)** Alignment of complete and largely complete *Mavericks* from nematode genomes. Highlighted in gray are homologous sequences with high nucleotide sequence identity. (*) The *C. bovis* scaffold is circular. **(C)** Alphafold2 prediction of *Cwal-*KRMA-1 protein structure. **(D)** Selected branches of the KRMA phylogeny. Full phylogeny containing 370 proteins is available in Figure S24. Colour code is the same as in (A).

Next, we asked whether *Mavericks* themselves were being horizontally transferred, regardless of their association to WASP proteases. To do so, we first annotated the core set of *Maverick* genes across 107 available nematode genomes and reconstructed their phylogeny (Figure S20). For all *Maverick* genes inspected, we found widespread incongruencies between gene and species trees, supporting multiple independent *Maverick* transfer events between species belonging to different genera, including *Caenorhabditis, Pristionchus, Panagrellus, Heterorhabditis, Mesorhabditis, Parapristionchus*, and *Micoletzkya* (Figure S20). This scenario is also supported by the extreme nucleotide identity of *Maverick* genes found in nematodes from different genera (Figure S21), as well as their GC content, which is lower than that of their host (Figure S22). Our analysis also revealed that most nematode *Mavericks* are either incomplete or pseudogenized, as is generally the case in eukaryotes [62]. However, we identified some noteworthy exceptions. For instance, *Mavericks* found in *C. wallacei, C. plicata, C. bovis, M. belari*, and *H. bacteriophora* were largely complete (Figure 5B). The *Mavericks* found in *C. wallacei, C. plicata*, and *C. bovis* were over 95% identical to each other at the nucleotide level, suggesting recent HGT events (Figure 5B). A common feature across *Mavericks* was the presence of a ∼2-5 kb region between the left TIR and the integrase, that could be distinguished due to its high GC%, low nucleotide conservation, and the presence of genes with introns, such as *Cpli-wasp-2* (Figure 5B). These sequences likely correspond to cargo genes that are taken up from the host and exchanged between species (Figure 5B; Figure S22).

### A novel family of kinases is also associated with Mavericks

Some of the largely complete *Mavericks* did not carry a *wasp* cargo but a gene of unknown origin. To characterize the new cargo, we first focused on the *C. wallacei Maverick*. An InterPro scan of its predicted 430 amino acid-encoding cargo revealed a ubiquitin-like fold at the N-terminus [AA:1-73], but no additional known domains. We then predicted its structure using AlphaFold2 (Figure 5C) and searched for structurally similar proteins in the PDB. The novel cargo was structurally homologous to numerous bacterial kinase effectors, virulence factors, and toxins including *Legionella pneumophila* MavQ, *Helicobacter pylori* CtkA, and *Shewanella oneidensis* HipA (top Z-score=11.9; Figure S23). Structural alignment of the new cargo to *H. pylori* Ser/Thr kinase CtkA revealed conservation of its key catalytic residues (RMSD=4.8Å; TM-score=0.59; Figure S23), suggesting that the *Maverick* cargo is a kinase [63]. In light of this result, we named the new cargo *Cwal-krma-1* (for *<u>K</u>inase-<u>R</u>elated family associated with <u>MA</u>vericks)*. To test whether KRMA kinases were being horizontally transferred, similar to WASP proteases, we comprehensively annotated KRMA kinases across 107 nematodes species and built a phylogenetic tree that included 532 proteins (Data S2). Similar to WASP proteases, we found widespread incongruencies in the gene tree compared to the species phylogeny and extreme DNA conservation in both coding and intronic regions between KRMA kinase homologs in highly divergent species (Figure 5D; full phylogeny in Figure S24). Furthermore, 22.7% (121 out of 532) of KRMA kinases were associated with either complete or partial *Mavericks* (Figure S24).

In summary, our results show that two nematode protein families—WASP proteases and KRMA kinases—are preferentially associated with *Mavericks* and as a result, broadly exchanged between species that have been reproductively isolated for tens or even hundreds of millions of years.

### Modularity of ubiquitin-like domains in cargo proteins

Besides being associated with *Mavericks*, WASP proteases and KRMA kinases share another feature. Both protein families have ubiquitin-like domains at their N-terminus despite sharing low levels of sequence identity with ubiquitin (Figure 3A; Figure 6A). The N-terminus of MSFT-1 contains three lysines (Lys8, Lys12, and Lys25) that are structurally analogous to lysines involved in ubiquitin chain formation, and a fourth lysine (Lys29), which is the most highly conserved of the four across WASP proteases (Figure 6B-C). To better understand the role of the ubiquitin-like domain of MSFT-1, we mutated the highly conserved Lys29 to alanine in the endogenous locus using CRISPR/Cas9. We observed no abnormal phenotypes in the parental Lys29Ala mutant line (98.3% wildtype, n=116/118). We then crossed *msft-1* K29A mutant hermaphrodites to AF16 males and inspected their F_2_ progeny. Similar to the *msft-1* wildtype situation, we observed ∼27% of affected F_2_ individuals (n=70/254; Figure 6D). However, these individuals were not developmentally delayed as in the wildtype situation but their phenotypes were much more severe; 92.9% of affected F_2_ progeny died as either embryos or early larvae (n=65/70; Figure 6D). In a total of 254 F_2_ progeny analyzed, only 4 AF16 homozygotes were phenotypically wildtype (Table S1). These results suggest that the ubiquitin-like domain of MSFT-1 regulates its abundance and that disrupting this domain increases MSFT-1 dosage in the early embryo and, as a result, its toxicity. Lastly, we found that these ubiquitin-like domains are highly modular within each protein family. For example, the ubiquitin-like domains of MSFT-1 and *Cpar*-WASP-2 were 92% identical at the protein level, whereas their protease domains were only 21% identical (Figure 6E). In contrast, the ubiquitin-like domains of MSFT-1 and *Mbel*-WASP-8 were 27% identical, whereas their protease domains were 72% identical (Figure 6E). This pattern was also evident between MSFT-1 and *Cpar*-WASP-1 (Figure 3E). KRMA kinases were also modular (Figure 6E). This unusual pattern of genetic variation is consistent with recombination between different *Mavericks*, analogous to viral recombination during infection and replication cycles [64].

**Figure 6.**
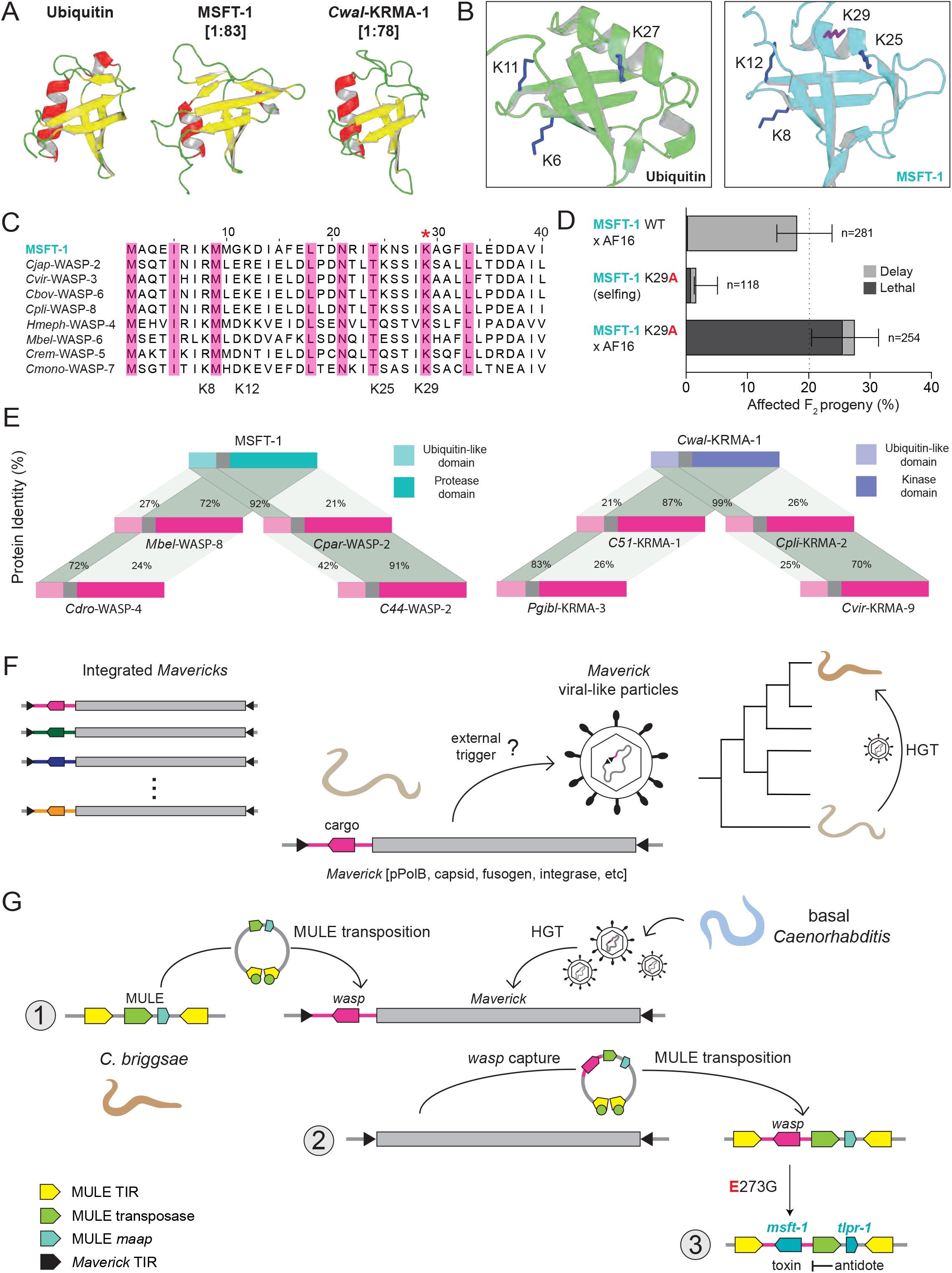
Modularity of *Maverick* cargo and model for Maverick-mediated HGT and the evolution of *msft-1/tlpr-1*. **(A)** Comparison of the structures of ubiquitin (PDB:1UBQ) and Alphafold2 predictions of the N-terminal domains of MSFT-1 and *Cwal*-KRMA-1. α-chains in red, β-sheets in yellow, linkers in green. **(B)** Structurally analogous lysines in ubiquitin and MSFT-1 ubiquitin-like domain (blue residues). A non-analogous lysine, K29, is present in MSFT-1 (purple residue). **(C)** Alignment of the N-terminal ubiquitin-like domain of MFST-1 and representative WASP proteases. K29 is the most conserved lysine (red asterisk). **(D)** A Lys29Ala mutation in MSFT-1 potentiates its toxicity in crosses, resulting in embryonic and larval lethality instead of developmental delay. Error bars indicate 95% confidence intervals calculated with the Agresti-Coull method. **(E)** Examples of modularity between the N-terminal ubiquitin-like domain and C-terminal catalytic domain of WASP proteases (left) and between the N-terminal ubiquitin-like domain and C-terminal catalytic domain of KRMA kinases (right). **(F)** Model for *Maverick*-mediated HGT of cargo genes in nematodes. We hypothesize that under some environmental perturbation, *Mavericks* are transcribed and form enveloped viral-like particles. These particles can infect individuals from other species and integrate into their germline. **(G)** Model for the origin of the *msft-1/tlpr-1* TA. (1) A *Maverick* from a basal *Caenorhabditis* species infected *C. briggsae* and integrated into the genome, introducing its cargo *wasp* protease (right). A *C. briggsae* MULE then transposed into this *Maverick* (left) and (2) captured the protease during subsequent transposition. (3) The *wasp* protease evolved into the *msft-1* toxin. The E273G mutation close to the catalytic site was critical for evolution of toxicity and may have changed the substrate specificity of the MSFT-1 protease domain. The *tlpr-1* antidote evolved from a *maap*, a small protein of unknown function associated with the MULE transposase, giving rise to the present-day MULE-associated *msft-1/tlpr-1* TA element in *C. briggsae*.

## Discussion

In this study, we provide evidence for widespread *Maverick*-mediated horizontal gene transfer across extremely divergent nematode species (Figure 6F) that drove the evolution of a novel MULE-associated selfish TA element, *msft-1/tlpr-1* (Figure 6G). The *msft-1/tlpr-1* TA is capable of both spreading in nature by poisoning non-carrier individuals and also changing its position in the genome via transposition. The existence of elements like *msft-1/tlpr-1* TA could explain why numerous transposable elements that are present in only one or two copies per genome can persist in the face of genetic drift [65]. Thanks to their association with TAs, transposons can rapidly spread in populations while incurring no fitness cost. Furthermore, elements like *msft-1/tlpr-1* provide a cautionary tale to evolutionary biologists, as they could be responsible for “ghost” signatures of selection in genomes. Quick fixation of the TA element followed by transposition could be erroneously interpreted as positive selection acting on neutral variants that were originally linked to the TA. The *msft-1/tlpr-1* TA is reminiscent of the *P*-element in *Drosophila* [2,11] and the *Spok* block, a recently reported selfish element in the fungus *Podospora anserina* [66]. The *Spok* block is made up of numerous *Spok* TAs capable of meiotic-drive, which are embedded within a putative giant DNA transposon, the *Enterprise*. Like *msft-1/tlpr-1*, the *Spok* block is a hyperparasite capable of both gene drive and transposition. However, there is a key difference between the two: *Spok* TAs can drive in populations independently of their association with transposable elements, whereas *msft-1/tlpr-1* cannot. The *msft-1/tlpr-1* TA is more than the sum of its parts. Its toxin evolved from a horizontally transferred protease, while its antidote was co-opted from a MULE-associated gene (Figure 6G). We show in unprecedented detail how cooperation between parasitic genetic elements can lead to the evolution of novel biological function, and ultimately impact gene flow between populations.

Our work reveals *Mavericks* as one of the long sought-after vectors of HGT. Thanks to their unique biology, which shares features of both transposons and viruses, *Mavericks* are responsible for the widespread transfer of genes across extremely divergent species. Analogous to endogenous retroelements that gained *env* genes, *Mavericks* captured a novel fusogen, MFUS-1, which is likely instrumental for HGT in nematodes [67,68]. However, we could not detect transcription of the fusogen or any other *Maverick* gene. We hypothesize that *Mavericks*—like *E. coli* λ-phage and human herpesvirus—integrate into the genome of their host and passively replicate until an environmental factor triggers the formation of infective particles (Figure 6F). Unlike *Mavericks* that are integrated in the genome of their host (Figure 5B), the *C. bovis Maverick* was found in a 17,366 bp scaffold that corresponds to a circular DNA sequence [69]. This *Maverick* has only one TIR, which connects the 3’ end of the pPOLb gene with the 3’ end of its cargo gene. This circular DNA could represent the genome of a virus-like particle after excision and recombination of the TIRs that flank the *Maverick*, or an intermediate state of transposition. Future studies focusing on the reconstitution of their life cycle will allow us to study HGT with unprecedented mechanistic detail *in vivo*. Moreover, this research could also lead to the development of payload delivery systems in nematodes, for which DNA viruses are unknown [70]. *Mavericks are* widespread in eukaryotes— pseudogenized copies are even present in humans—and their gene content is highly conserved [57,62,71]. Intriguingly, teleosts, such as ray-finned fishes, have experienced extreme levels of HGT and have a high number of *Mavericks* in their genomes [7,62]. Thus, we predict that *Mavericks*—and analogous selfish genes with viral properties—could mediate HGT in other animal lineages including vertebrates.

## Acknowledgments

We thank E. Haag, S. Baird, and C. Braendle for strains. We thank J. Ross and K. Senti for discussions, and P. Duchek, B. Érdi and J.Gokcezade from the IMBA Fly & Worm facility for the design and generation of transgenic worms. Library preparation and sequencing were performed at the VBCF NGS Unit. We thank Life Science Editors for editing services. Research in the Burga lab is supported by the Austrian Academy of Sciences, the city of Vienna, and a European Research Council (ERC) Starting Grant under the European Union’s Horizon 2020 program (ERC-2019-StG-851470). S.A.W. is supported by the European Union’s Framework Programme for Research and Innovation Horizon 2020 (2014-2020) under the Marie Curie Skłodowska Grant Agreement Nr. 847548. I.C.B. is supported by “la Caixa” Foundation (LCF/BQ/EU19/11710050). A.K. is supported by a Boehringer Ingelheim Fonds (BIF) PhD Fellowship. Sequencing data are available under NCBI project PRJNA857754.

## Material and Methods

### Maintenance of worm strains

All worms were maintained at 25°C on modified nematode growth medium (NGM) plates unless otherwise indicated. The modified NGM contains 1% agar/0.7% agarose as *C. briggsae* and other species can burrow into normal NGM plates. Some of the strains were provided by the CGC, which is funded by the NIH Office of Research Infrastructure Programs (P40 OD010440). All strains used in this study are listed in Table S3.

### Generation of *C. briggsae* near-isogenic and mutant lines

All near-isogenic lines were generated as previously described [23] and an overview is shown in Figure S1A. For CRISPR/Cas9 genome editing, we adapted previously described protocols [72]. In brief, 250 ng/µl Cas9 or Cas12a (IDT) protein was incubated with 200 ng/µl crRNA (IDT) and 333 ng/µl tracrRNA (IDT) at 37°C for 10 mins before adding 2.5 ng/µl co-injection marker plasmid (pCFJ90-mScarlet-I). For homology-directed repair (HDR), donor oligos (IDT) or biotinylated and melted PCR products were added at a final concentration of 200 ng/µl or 100 ng/µl, respectively. Following injections into young adult hermaphrodites, mScarlet-positive F_1_ progeny were singled and their offspring screened by PCR and Sanger sequencing to detect successful editing events. All gRNAs and HDR templates are listed in Table S4.

### Phenotyping and genotyping of crosses

All crosses were performed at 25°C. Crosses were set up in a single well of a 12-well plate with 6 L4-stage hermaphrodites and at least 35 adult males, then allowed to mate and lay for 48 hrs. For F_1_-selfing experiments, L4-stage F_1_s were then singled onto 3 cm plates and allowed to self-fertilize overnight. To ensure that the F_2_s used for phenotyping were approximately the same age, the F_1_ mothers were transferred to a new plate in the morning, allowed to lay for 2 hrs, then fresh F_2_ eggs were singled onto 3 cm plates for phenotyping. All F_2_s were then staged every 24 hrs to track their development. Eggs that did not hatch were scored as embryonic lethal, and larvae that hatched but arrested before sexual maturity were scored as larval lethal. At 48 hrs, wildtype worms have reached sexual maturity and are laying eggs; worms were considered to be developmentally delayed if they did eventually reach sexual maturity, but had not yet done so at 48 hrs (in most cases, delayed worms were L4-stage or just molted to adult-stage). In all cases, F_2_s were monitored until they either died or reached sexual maturity. All crosses were performed at least twice, with 10-20 F_2_ progeny analyzed from 5-10 different F_1_ mothers in each experiment. Each n-value given indicates the total number of progeny that were counted.

For maternal backcrosses, crosses were performed as above except that L4-stage F_1_s were transferred to a new 3 cm plate and mated with at least 50 males of the appropriate parental strain overnight, then singled in the morning to lay fresh eggs for analysis. Paternal backcrosses were performed by transferring at least 50 F_1_ young males to a new 3 cm plate, left overnight to ensure they reached sexual maturity, then at least 10 L4-stage hermaphrodites of the appropriate parental strain were added. They were allowed to mate overnight, then the hermaphrodites were singled in the morning and fresh eggs were picked for analysis.

Genotyping PCRs were performed with Proteinase K (ProK) digested worm lysates as template using in-house hot-start Taq polymerase with an annealing temperature of 56°C. ProK digestion was performed for 30 mins at 37°C with 200 ug/mL enzyme diluted in Taq buffer, then 1 uL was used directly in genotyping PCRs. In all crosses where we observed F_2_ delay, all F_2_ progeny were genotyped regardless of their phenotype, by PCR using template from ProK digestion of either the F_2_ worm itself or at least 10 pooled F_3_s. The genotypes for all relevant crosses can be found in Table S1. In crosses where we observed no delay, all F_1_ mothers and at least 40 F_2_s were genotyped from at least two different confirmed heterozygous F_1_ mothers to ensure that we observed the expected Mendelian ratios of all genotypes. Where applicable, the genotyping of all crosses and lines used in this study were performed with either AFLP assays or Sanger sequencing. A list of all primers and genotyping assays can be found in Table S5.

### Short-read Illumina whole-genome sequencing

We extracted genomic DNA (gDNA) from at least two freshly starved 9 cm plates of the appropriate worm strain using the MasterPure Complete DNA and RNA Purification kit (Lucigen; cat. #MC85200), following manufacturer protocols, including the sample freeze/thaw step to facilitate tissue lysis. We then prepared Illumina sequencing libraries using the Nextera DNA Flex Library Prep kit (Illumina; cat. #20018705) with 100 ng of input gDNA for each library. gDNA and prepared libraries were quantified using the DeNovix dsDNA High Sensitivity kit (Biozym; cat. #31DSDNA-HI1), libraries were checked on a Fragment Analyzer and then sequenced on an Illumina NextSeq or MiSeq instruments. All sequencing runs used in this study are listed in Table S6.

### RNA extraction and RNA-seq

For AF16 and HK104 mixed-stage embryo RNA-seq, three full, non-starved 9 cm plates of the appropriate worm strain were first bleached to isolate only the embryos, then total RNA was extracted from them using a TRIzol extraction method based on a previously published protocol [73] with the following modifications - chloroform was used instead of bromochloropropane, and phase-lock tubes were used during TRIzol/chloroform steps. DNA was removed from the samples with a DNase I digestion (30 mins at 37°C), then the enzyme was inactivated with 2.5 µL of 25 mM EDTA and a 10 mins incubation at 65°C. 100 ng of purified RNA was used as input for the TruSeq Stranded mRNA kit (Illumina; cat. #20020595) and libraries were prepared according to manufacturer instructions. Libraries were then checked on a Fragment Analyzer and sequenced on a 2×75 paired-end NextSeq550 lane. All samples were run in biological triplicate.

### Single-molecule Nanopore sequencing

High molecular weight DNA was extracted using either the MasterPure Complete DNA and RNA Purification kit (Lucigen; cat. #MC85200), or by standard phenol/chloroform extraction. Worms were washed off of freshly starved plates and rinsed with M9 at least 3 times before the pellet was frozen at -20 to facilitate sample lysis before the extraction protocol was started. Either 10 kb (*C. briggsae* strains) or unfragmented (*C. plicata* SB355) sequencing libraries were prepared using the 1D Ligation Sequencing Kit from Oxford Nanopore Technologies (ONT; SQK-LSK109) and sequenced on a PromethION device (ONT). Libraries for *C. briggsae* HK104 and *C. plicata* SB355 were run individually, while *C. briggsae* ED3036, JU439 and QR24 were barcoded using the Native Barcoding kit (#EXP-NBD104) and sequenced together. Read calling and demultiplexing (where applicable) was performed with the GPU-version of Guppy Basecalling software v5.0.11+2b6dbffa5 and the basecalling model dna_r9.4.1_450bps_sup_prom.cfg.

### Genome assembly and annotation

Raw Nanopore reads were error corrected using NECAT v0.0.1 with an estimated genome size of 105 Mb. The longest corrected reads were taken so that the estimated coverage of the whole genome would be 50x. These reads were then trimmed by Canu v1.8 and assembled using Flye v2.7.1 with default parameters [74,75]. The resulting assembly was polished by two to four rounds of Racon v1.4.2. Scaffolds corresponding to residual *E. coli* contamination were identified and removed using BLAST v2.2.26. For the *C. plicata* assembly, Flye was run in the metagenomic mode due to extensive bacterial contamination of the library. Circular contigs with coverage more than twice higher than coverage of true *C. plicata* contigs were removed. True contigs were defined as nearly chromosome-sized contigs showing sequence homology to the existing *C. plicata* short-read assembly. Genome annotation was carried out using funannotate v1.7.4, with a transcriptome assembled by Trinity v2.10.0 [76,77]. For funannotate, Augustus was run with a pre-trained *C. elegans* model, BUSCO was run with -- busco_db nematoda --busco_seed_species caenorhabditis options. GeneMark-ET was run with default parameters.

### Fine mapping of the toxin-antidote element in HK104 Chr. III

We began mapping the HK104 TA by performing 16 generations of maternal backcrosses to the non-carrier strain AF16, taking advantage of the ability of TAs to select for themselves in each generation. For each backcross, 5-10 L4-stage hermaphrodites were crossed with at least 25 AF16 males in a 3 cm plate. The 16th generation was Illumina short-read sequenced (see “Short-read Illumina whole-genome sequencing” section; Table S6) and we performed SNP-calling analysis to calculate allele frequencies across the genome (see “SNP-calling analysis” section). This allowed us to identify a single large introgression from HK104 on Chr. III (approximately 0-10 Mb; Figure 1B) in an otherwise AF16 background. We then isolated 50 independent near-isogenic lines (NILs) from the 16x backcrossed generation by picking individual L4-stage hermaphrodites and allowing them to self-fertilize. The introgressions carried by each NIL were roughly mapped using a combination of previously published [27] and newly identified marker indels (Table S5) across Chr. III that had large enough size difference in the AF16 and HK104 genomes to be visualized by PCR and gel electrophoresis. We identified two particularly useful NILs (NIL1 and NIL2) that had partially overlapping introgressions but both carried the HK104 TA (Figure S1A-C). The exact margins of the overlap between NIL1 and NIL2 were determined by Illumina short-read sequencing, alignment of each dataset to the AF16 genome, and manual inspection of SNPs using IGV (Chr. III: 4.94-6.75 Mb in AF16 coordinates). NIL1 was used for all further experiments in the paper and carries a 3.24 Mb introgression (Chr. III: 3.51-6.75 Mb; Figure S1B). To further narrow the candidate region, we analyzed existing AF16/HK104 recombinant interbred lines (RILs) that had different recombination breakpoints across the introgression (Figure 1C) by genotyping them for the same Chr. III markers and also crossing them to AF16 and analyzing their F_2_ progeny for developmental delay to determine if they carried the HK104 TA. We then used our HK104 *de novo* assembled genome (see “Genome assembly and annotation” section) to establish a list of predicted genes within the overlap region, and narrowed the list of candidates first based on their gene expression during embryonic development using our RNA-seq data from both HK104 and AF16 (see “RNA extraction and RNA-seq” section). After filtering for genes that were expressed in HK104 and not expressed in AF16 (or, if expressed, highly divergent), we then searched for pairs of candidates that were immediate neighbors or in tight linkage to each other (Table S2). Only one pair of genes, *ORF14767* and *ORF14768* satisfied all criteria, and they were subsequently targeted with CRISPRs to generate knockout alleles (see “Generation of *C. briggsae* near-isogenic and mutant lines” section). *ORF14767* mutants and *ORF14767 ORF14768* double mutants were tested in genetic crosses as described previously [23] to confirm that they are the toxin-and antidote-encoding genes, respectively. All mutant lines are available in detail in Table S3.

During analysis of the antidote, we identified a highly conserved copy of *ORF14768* on Chr. V in multiple *C. briggsae* wild isolates and the reference strain AF16 (*CBG17069*.*1*). The predicted coding sequences of *ORF14768* and *CBG17069*.*1* have 91.8% identity; their predicted proteins only differ in the first 14 amino acids and three discrete substitutions later in the peptide (p.K95E, p.C99S, p.F151L). We used genetic crosses and genotyping to rule out the possibility that *CBG17069*.*1* functions as an antidote against the HK104 toxin. Due to its high sequence similarity to *ORF14768, CBG17069*.*1* was also targeted by the sgRNAs when we generated the *ORF14768* HDR allele (see “Generation of *C. briggsae* near-isogenic and mutant lines” section). Consistent with our prediction, we saw no correlation between the F_2_ delay phenotype and *CBG17069*.*1* genotype. This is also consistent with AF16 being susceptible to the *msft-1/tlpr-1* TA in crosses, despite evidence that *CBG17069*.*1* is expressed in AF16 [78]. Together, these data support the conclusion that *ORF14768* is the cognate antidote to *ORF14767*, and that *CBG17069*.*1* is not, despite the sequence similarity. To specifically genotype *CBG17069*.*1* we used the same PCR conditions described above (see “Phenotyping and genotyping of crosses” section) (F primer: 5’-tcccaagttcagtgattttcct, R primer: 5’-tctcgtactcaaacccaatcct) followed by Sanger sequencing with the reverse primer, using two SNPs in exon 3 to differentiate between *ORF14768* and *CBG17069*.*1* (c.A283G, c.T295A).

### SNP-calling analysis

The reads were aligned to the reference genome using SAMtools v1.10. Duplicates were removed using sambamba v0.6.6. Variants were called using GATK v4.0.1.2. Variants were quality filtered with a threshold of QUAL=200. Fractions of reads supporting one or another allele were estimated using custom R script. All scripts are available upon request.

### Annotation of MULE transposases and phylogenetic reconstruction

For the annotation of MULE catalytic transposases we used OrthoFinder v2.5.4 with default settings [79]. As an input, we used a combination of genomes generated in this study and those available in two public databases: *Caenorhabditis* Genomes Project (V1) and Wormbase Parasite (WBPS16). After obtaining the MULE transposase orthogroup we exclusively kept the MULE catalytic domains and sequences below 100 amino acids were removed. After the filterings we obtained 161 MULE catalytic transposase sequences. Then, the sequences were aligned with MAFFT v.7.427 with default options [80]. With the purpose of removing spurious sequences we used trimAL [81] with the following options (*-resoverlap 0*.*75 -seqoverlap 80*). To build the phylogeny we used IQ-TREE v.1.6.12 [82] with the following parameters (*-st AA -m TESTNEW -bb 1000 -alrt 1000)*. The phylogenetic tree was coloured using the interactive Tree Of Life (iTOL software version 6.5.7) [83].

### *De novo* protein structure prediction and analyses

We predicted protein structures using AlphaFold2 [34] and RoseTTAfold. AlphaFold2 was run using the ColabFold *AlphaFold2_advanced* notebook implementation in Google Colaboratory [84]. For multiple sequence alignment, we selected the mmseqs2 option. Models were ranked based on pLDDT and only the best ranked out of five models was selected for further analysis. We searched for structurally homologous proteins on the protein database base (PDB) using the DALI server [36]. Graphics were generated using PyMol (The PyMOL Molecular Graphics System, Version 2.5 Schrödinger, LLC.) Protein alignment statistics for selected pairs were generated with TM-align [85] and PyMol using the *super* algorithm for protein pairs with high sequence identity and *cealign* for protein pairs with low sequence identity. We estimated and plotted the evolutionary conservation of MSFT-1 residues using the ConSurf server [86].

### Horizontal gene transfer analysis of *msft-1*

We first obtained genome-wide single-copy orthologs from *C. briggsae HK104, C. plicata, C. parvicauda*, and *C. auriculariae* using OrthoFinder v2.5.4 [79]. We selected these three additional species because they carried proteases that were most closely related to *msft-1*. The sequences for each pair of orthologs were codon-aligned using the MACSE v2.04 software [87]. We then used the VHICA (Vertical and Horizontal Inheritance Consistence Analysis) method to discriminate between vertical and horizontal gene transfer which is implemented in the *vhica* R package [54].

### Annotation of WASP proteases and KRMA kinases and phylogenetic reconstruction

For the annotation of WASP proteases and KRMA kinases protein families we used OrthoFinder v2.5.4 with default settings [79]. As an input, we used a combination of genomes generated in this study and those available in two public databases: *Caenorhabditis* Genomes Project (V1) and Wormbase Parasite (WBPS16). Proteins identified were randomly assigned a WASP and KRMA number in each species irrespective of orthology between species. Orthogroups for each family were then aligned with MAFFT v.7.427 with default options [80]. In order to increase the phylogenetic signal from our protein alignment, we used trimAl v.1.4.1 to remove spurious sequences from the input alignment [81]. trimAL was run with the following options for the WASP family (*-resoverlap 0*.*75 -seqoverlap 80*). Given the high divergence of the KRMA family, we used a gap threshold of 0.5 with the option *-gt* (gap threshold is the minimum fraction of sequences without a gap). Then we used IQ-TREE v.1.6.12 to build the protein phylogenetic tree with the following parameters: *-st AA -m TESTNEW -bb 1000 -alrt 1000)* [82,88,89]. The phylogenies were coloured using the interactive Tree Of Life (iTOL) software version 6.5.7 [83].

### Annotation of *Mavericks* and phylogenetic reconstruction

We used tblastn (blast+ v. 2.8.1) to identify highly conserved Maverick genes in the genome of all nematode species used in this study [90]. We used as a query the following genes: ATPase, Fusogen, Integrase, Major capsid, minor capsid and PolB. We filtered out tblastn hits with an e-value above 0.0001 and those genes that were shorter (<60%) than the consensus *Maverick* gene (except for pPolB Maverick genes, sequences below 500 bp length were filtered out). We aligned sequences with MUSCLE v.3.8.31 (default options) and IQ-TREE v.1.6.12 was used to build the protein phylogenetic tree [82,88,89].

### Genomic association between *Mavericks* and *wasp* and *krma* gene families

We extracted the *krma* and *wasp* genomic coordinates for every sequence found in each of the two orthogroups. We did the same for the Maverick genes annotated in the previous step. We made a call for association when *wasp* and *krma* genes were 6 kb up and/or downstream from a *Maverick* gene.

### Comparison of GC content between Mavericks and host genes

We concatenated all core-genes (CDS) and all Maverick genes of the following species: *C. briggsae* HK104, *C. plicata, C. parvicaud*a and *C. auriculariae*. We used the SeqKit software to estimate the GC content of Maverick and core-genes [91]. To show the different GC content of Mavericks with respect to the rest of the genome, we estimated the GC content in a sliding window manner in two nematodes (*C. auriculariae* and *O. tipula*e) in a genomic region (∼500 kb) in which a Maverick is located.

### Genomic alignments

Sequence alignments of genomic regions containing MULEs and *Mavericks* were generated using Minimap2 [92]. We obtained pairwise sequence alignments using the following parameters *-X -N 50 -p 0*.*1 -c*. Final plots were generated using the *gggenomes* R package (https://github.com/thackl/gggenomes).

