## Supplementary Figures and Tables for "Virus-like transposons cross the species barrier and drive the evolution of genetic incompatibilities"

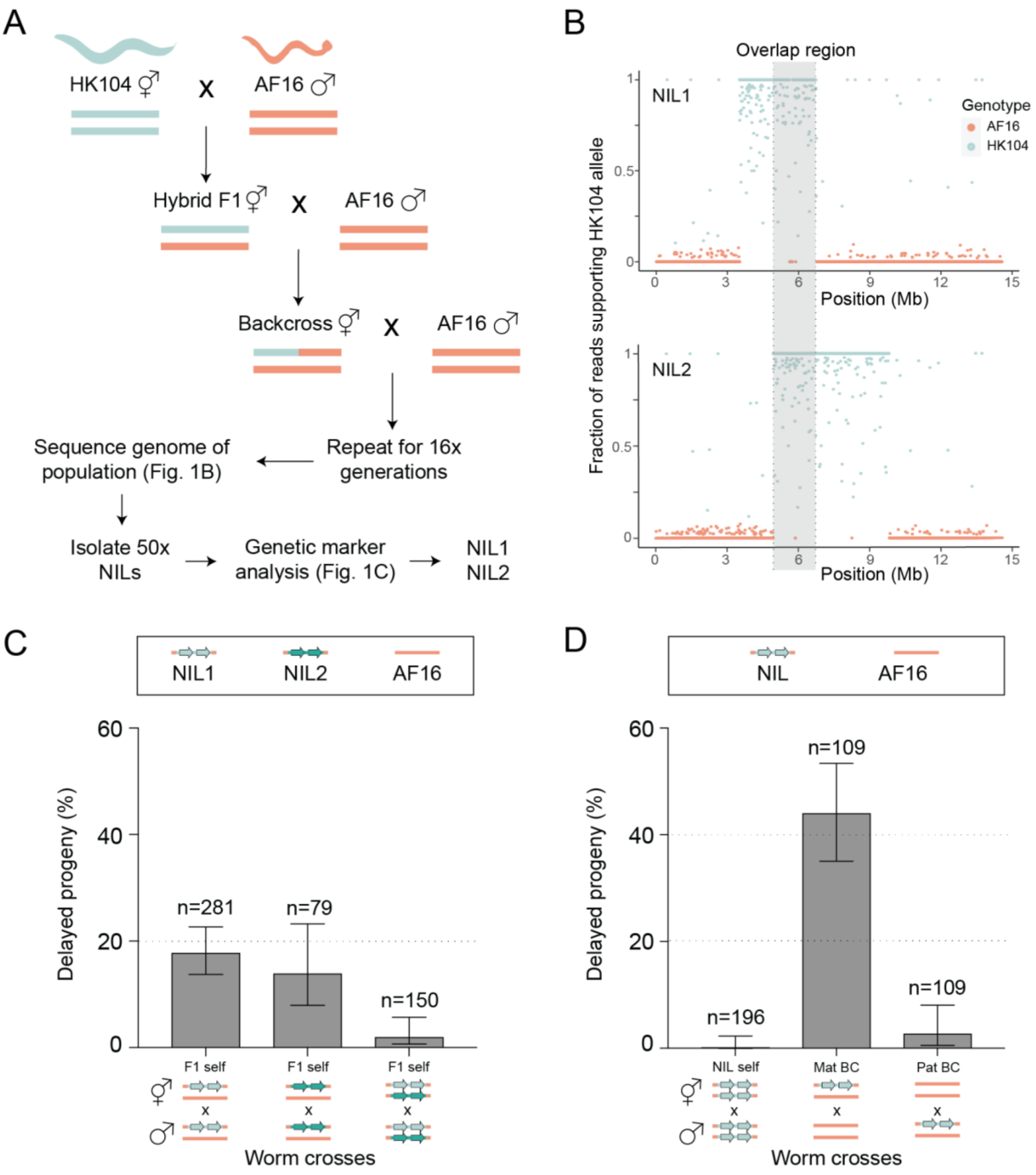


**Figure S1. Generation and genetic characterization of near-isogenic lines. (A)** Outline of backcrossing scheme to generate HK104>AF16 near-isogenic lines (NILs). The HK104 toxin acts through the maternal germline, so hybrid hermaphrodites were backcrossed to AF16 males for 16 generations. The 16x backcross genomes were Illumina short-read sequenced to identify regions of the HK104 genome that were retained. We isolated 50 individual L4 hermaphrodites and established NILs, then analyzed the genotype at multiple genetic markers across the introgression. We chose two NILs with partially overlapping introgressions for further analysis. **(B)** Allele frequencies across Chr. III in NIL1 (top) and NIL2 (bottom). The overlap region between their introgressions (illustrated by fixed HK104 alleles in cyan) is highlighted in gray. **(C)** Test crosses with NIL1 and NIL2. Both NILs were crossed to the non-carrier strain AF16; in both cases, we observed the F2 delay phenotype, indicating that both NIL1 and NIL2 carry the HK104 TA element. As a further control, we also crossed the NILs together and observed no F2 delay, consistent with both parents sharing the TA element. All further experiments used NIL1, now simply referred to as “NIL”. **(D)** Test crosses with the NIL. As expected, when the NIL is crossed to AF16, the rate of delay in maternal backcross (BC) progeny doubles (compare to NIL1 F1 self in C), while only background levels of delay are seen in the paternal BC. This is consistent with a maternal-acting toxin and recapitulates the results of crosses with the parental HK104 to AF16 (Ben-David et al. 2021).

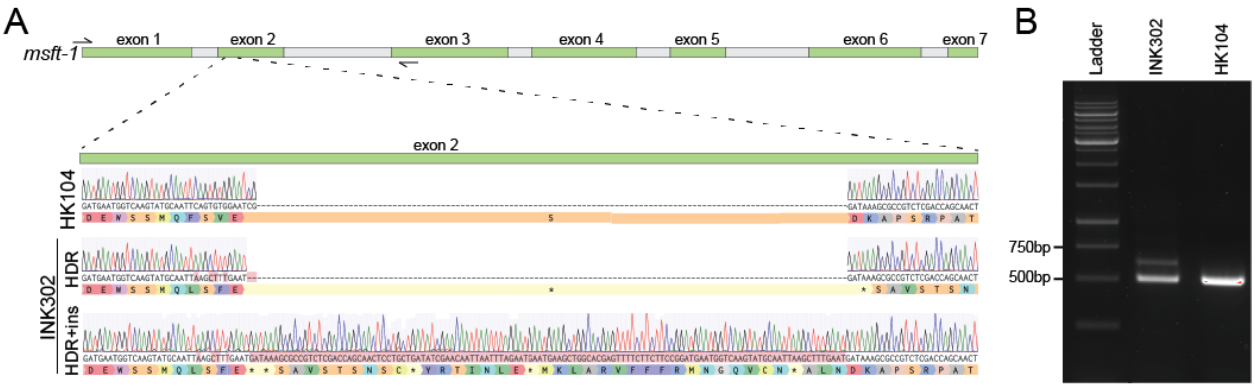


C

```
                  1                                                                                82
MSFT-1 wildtype   MAQEIRIKMMGKDIAFELTDNRITKNSIKAGFLLEDDAVISLSYEKNAHQICCQMNEAGTSFLLPDEWSSMQFSVESDKAPS
MSFT-1 HDR        MAQEIRIKMMGKDIAFELTDNRITKNSIKAGFLLEDDAVISLSYEKNAHQICCQMNEAGTSFLLPDEWSSMQLSFE------
MSFT-1 HDR+ins    MAQEIRIKMMGKDIAFELTDNRITKNSIKAGFLLEDDAVISLSYEKNAHQICCQMNEAGTSFLLPDEWSSMQLSFE------

                  ..................................................................................

                  83                                                                              164
MSFT-1 wildtype   RPATPADIEGPNPKKRKLDPYPEQLAVNGDLAASIGRYALYYNVKRDNYKRCAIPLTPRLAATFHHGENKSIKIGDEIVIHS
MSFT-1 HDR        ----------------------------------------------------------------------------------
MSFT-1 HDR+ins    ----------------------------------------------------------------------------------

                  ..................................................................................

                  165                                                                             246
MSFT-1 wildtype   WLDKNLKVATRAVKIMEEYDTIILQADGEHLSDKDFITSSGYPQKGMKYLLMGFSIIHEKTSHLSLSSGIIATEVTRRLRYT
MSFT-1 HDR        ----------------------------------------------------------------------------------
MSFT-1 HDR+ins    ----------------------------------------------------------------------------------

                  ..................................................................................

                  247                                                                          325
MSFT-1 wildtype   GSSGSFKGDSGGGCWSEDGHLMGMQIGVEKVPHTKDTNGRPASPASGGRCCIIAIRDILGHIQDLMPMPADSDADSWSE
MSFT-1 HDR        -------------------------------------------------------------------------------
MSFT-1 HDR+ins    -------------------------------------------------------------------------------

                  ...............................................................................
```

**Figure S2. Characterization of toxin null allele. (A)** The *msft-1* gene structure and Sanger sequencing of wildtype (HK104) and mutant alleles in INK302. During creation of the mutants, *msft-1* was duplicated and both copies carry a different mutant allele - a 2bp deletion that introduces a premature stop codon (HDR), and the 2bp deletion followed by a 102bp insertion (HDR+ins). Exons are shown in green, introns are shown in gray. **(B)** Gel electrophoresis of the genotyping amplicon (primers indicated by half arrows in exon 1 and 3 in A) showing both mutant band sizes compared to HK104 wildtype control. This image is cropped and the source gel is available in supplemental material. **(C)** Protein alignment of wildtype and mutant MSFT-1. Both mutant alleles in INK302 produce the same MSFT-1 peptide that is truncated after p.E76.

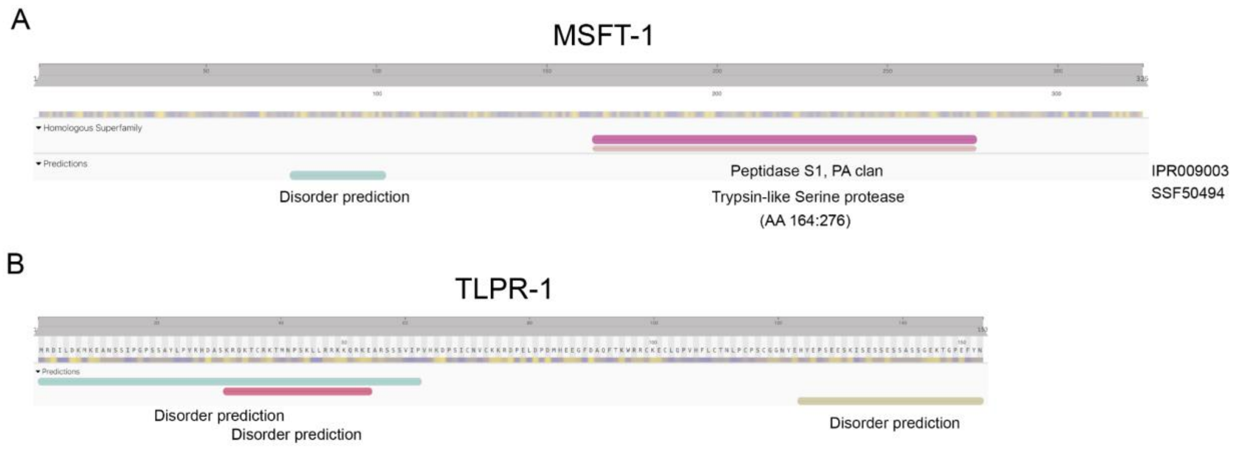


**Figure S3. InterPro domain predictions for the toxin and its antidote. (A)** InterPro domain prediction for the MSFT-1 toxin. A trypsin-like serine protease domain is predicted on the C-terminal region (AA 164:276). A disordered region is predicted in the linker region. **(B)** InterPro domain prediction for the TLPR-1 antidote. No known protein domains are detected. Both the N-terminus and C-terminus are predicted to be disordered.

A

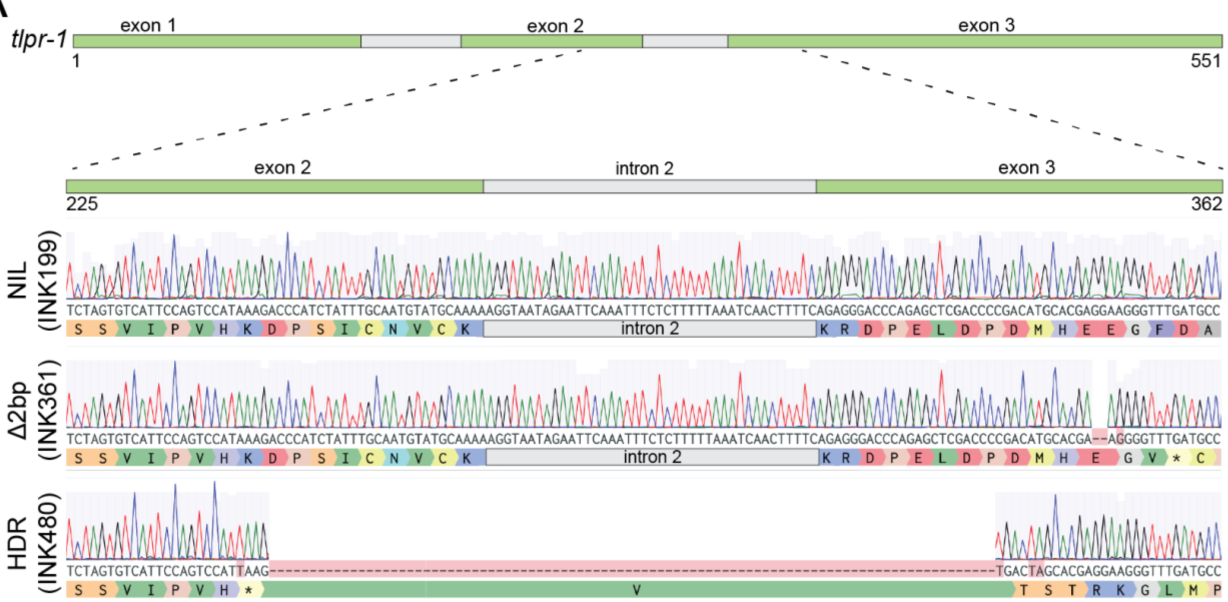


B

```
                  1                                                                                82
TLPR-1 wildtype   MRDILDKMKEANSSIPGPSSAYLPVRHDASKRQKTCRKTMNPSKLLRRKKQRKEARSSSVIPVHKDPSICNVCKKRDPELDP
TLPR-1 Δ2bp       MRDILDKMKEANSSIPGPSSAYLPVRHDASKRQKTCRKTMNPSKLLRRKKQRKEARSSSVIPVH------------------
TLPR-1 HDR        MRDILDKMKEANSSIPGPSSAYLPVRHDASKRQKTCRKTMNPSKLLRRKKQRKEARSSSVIPVHKDPSICNVCKKRDPELDP
                  ..................................................................................

                  83                                                                 153
TLPR-1 wildtype   DMHEEGFDAQFTKWRRCKECLQPVHFLCTNLPCPSCGGNYEHYEPSEESKISESSESSASSGEKTGPEFYN
TLPR-1 Δ2bp       -----------------------------------------------------------------------
TLPR-1 HDR        DMHEGV-----------------------------------------------------------------
                  .......................................................................
```

**Figure S4. Characterization of antidote null alleles. (A)** The *tlpr-1* gene structure and Sanger sequencing of wildtype and mutant alleles. We generated two different antidote mutant lines, one carrying a 2bp deletion that results in a frameshift and premature stop codon, and one carrying an HDR repair sequence that introduces a premature stop codon followed by a 90bp deletion. Exons are shown in green, introns are shown in gray. **(B)** Protein alignment of wildtype and mutant TLPR-1. Both *tlpr-1* alleles result in a truncated protein, p.E87GfsX2 and p.K65X, respectively. Both mutations were made in the *msft-1* mutant background (lines are *msft-1 tlpr-1* double mutants).

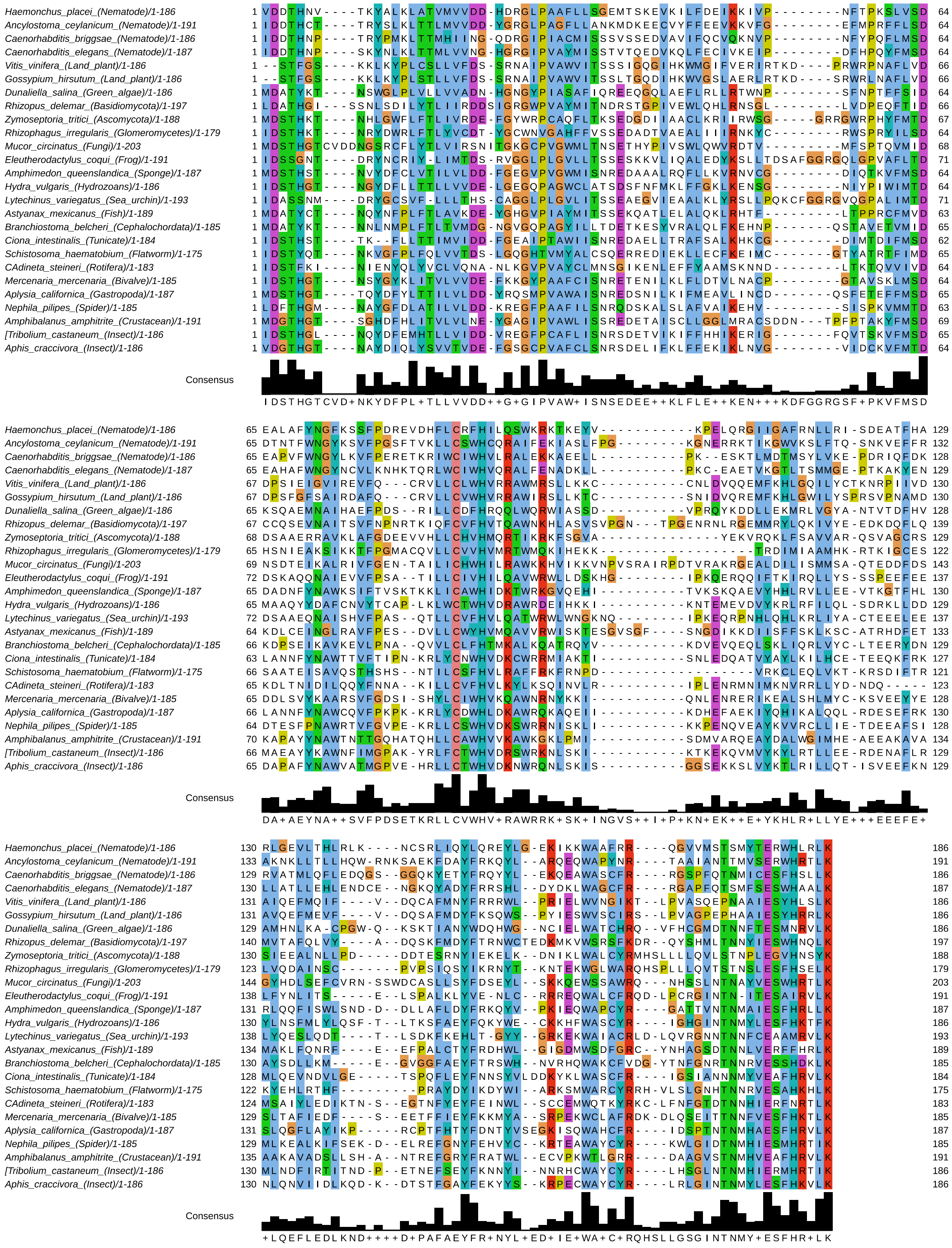


**Figure S5. Conservation of the *C. briggsae* MULE transposase across eukaryotes.** MULE-catalytic domain alignment covering 26 eukaryote species. Sequences from non-nematode species were retrieved from NCBI by running PSI-BLAST (the *C. briggsae* HK104 MULE-catalytic domain was used as the query sequence).

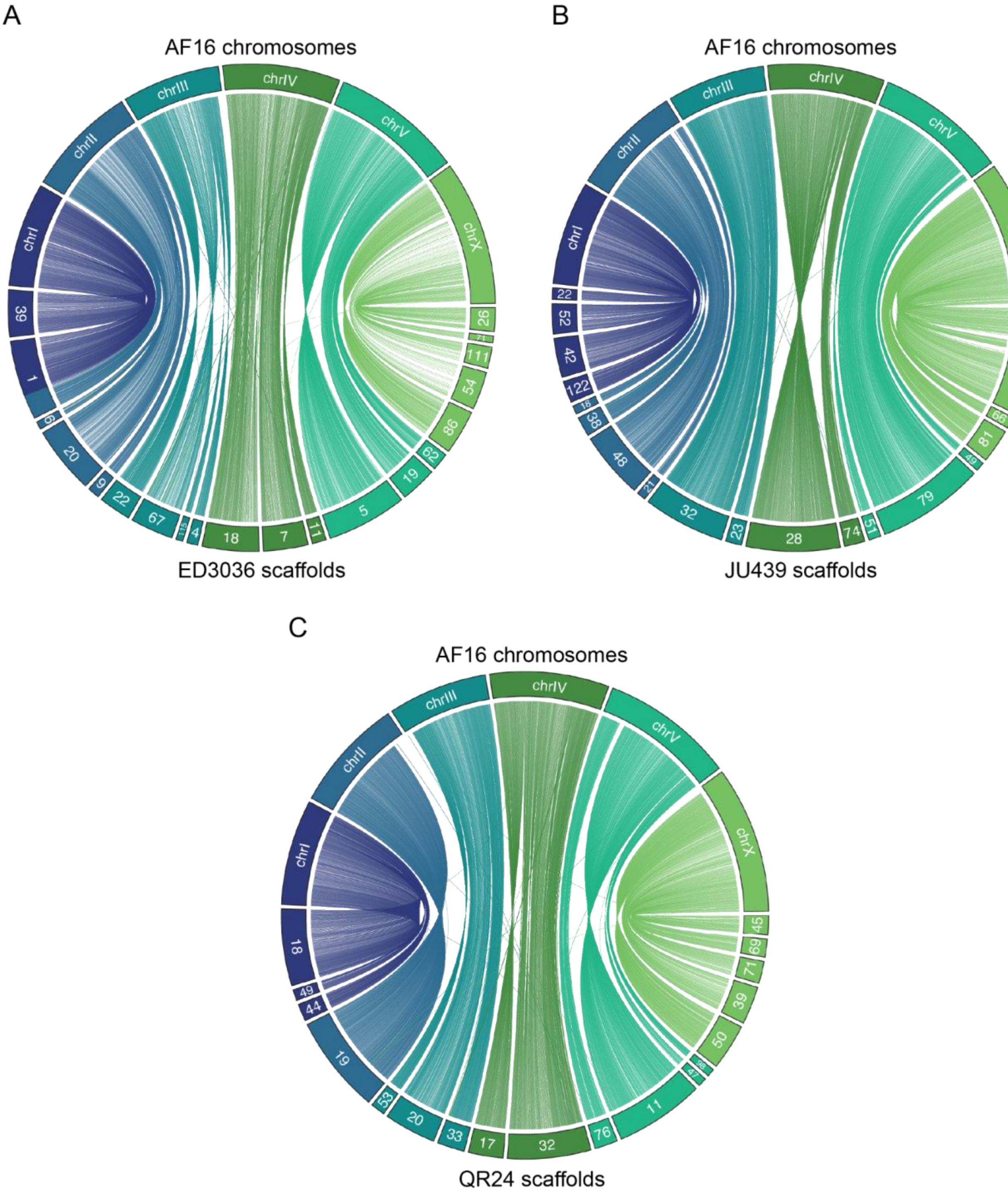


**Figure S6. Genome assembly of additional *C. briggsae* wild isolates. (A-C)** Synteny analysis between *C. briggsae* reference strain AF16 chromosomes and *de novo* assembly scaffolds for *C. briggsae* wild isolates ED3036 (A), JU439 (B), and QR24 (C). Only scaffolds above 1Mb are shown.

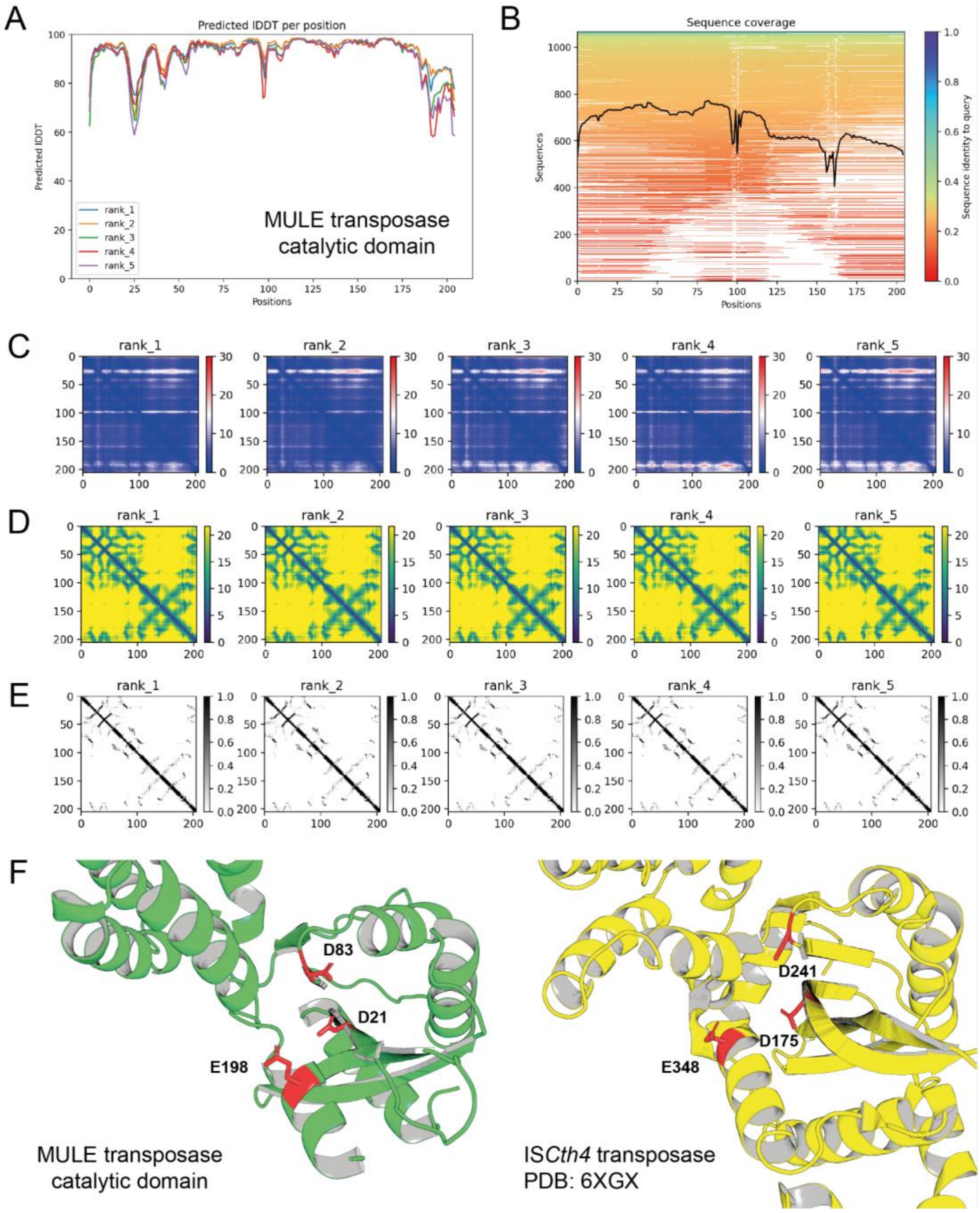


**Figure S7. Structural homology between the *C. briggsae* MULE transposase and the bacterial IS*Cth4* transposase. (A)** AlphaFold2 per-residue confidence metric pLDDT (Local Distance Difference Test). **(B)** Coverage of the multiple sequence alignment. **(C)** Predicted aligned error, confidence in the domain packing and large-scale topology of the protein. **(D)** Predicted distogram. **(E)** Predicted contacts. **(F)** Comparison of the core transposase domain of the *C. briggsae* MULE linked to *msft-1* and IS*Cth4,* a bacterial transposase of the *Mutator* family (PDB: 6XGX). Highlighted in red are the three DDE catalytic residues of IS*Cth4* and those identified in the MULE transposase as the catalytic triad based on structural homology and evolutionary conservation.

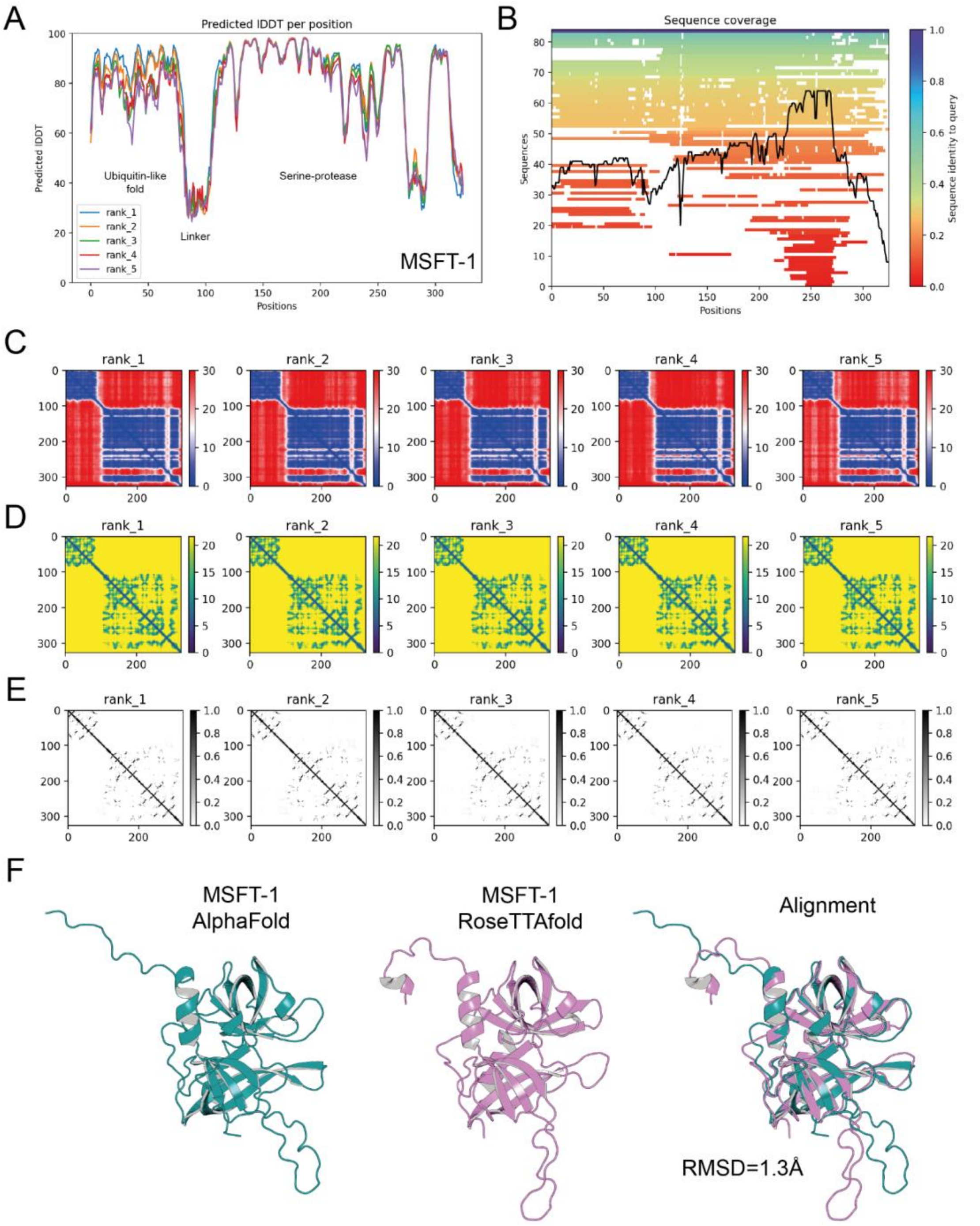


**Figure S8. Structural prediction of MSFT-1 toxin. (A)** AlphaFold2 per-residue confidence metric pLDDT (Local Distance Difference Test). The linker region in between the N-terminal ubiquitin-like fold and the C-terminal protease is largely unstructured. **(B)** Coverage of the multiple sequence alignment. **(C)** Predicted aligned error, confidence in the domain packing and large-scale topology of the protein. **(D)** Predicted distogram. **(E)** Predicted contacts. **(F)** Comparison of MSFT-1 models generated by Alphafold2 and RoseTTAfold for the protease domain. Both structures are highly similar except for an unstructured loop towards the end of the protein.

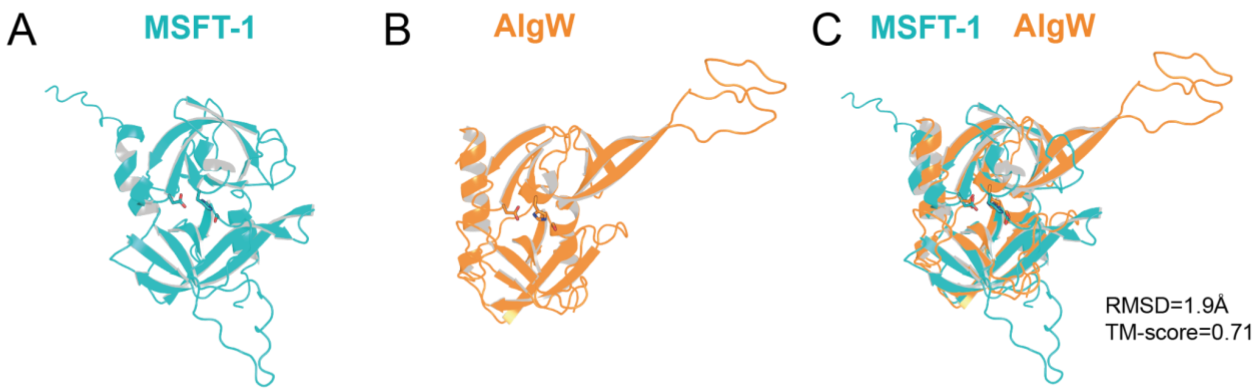


**Figure S9. Structural alignment of AlgW and MSFT-1 proteases. (A)** Alphafold2 prediction of the serine-protease domain of *C. briggsae* MSFT-1. **(B)** Structure of the HtrA-Type protease AlgW from *Pseudomonas aeruginosa* generated by X-ray diffraction (PDB: 7CO5). **(C)** Structural alignment between the protease domain of MSFT-1 and AlgW. The side chains of the catalytic residues are shown as sticks.

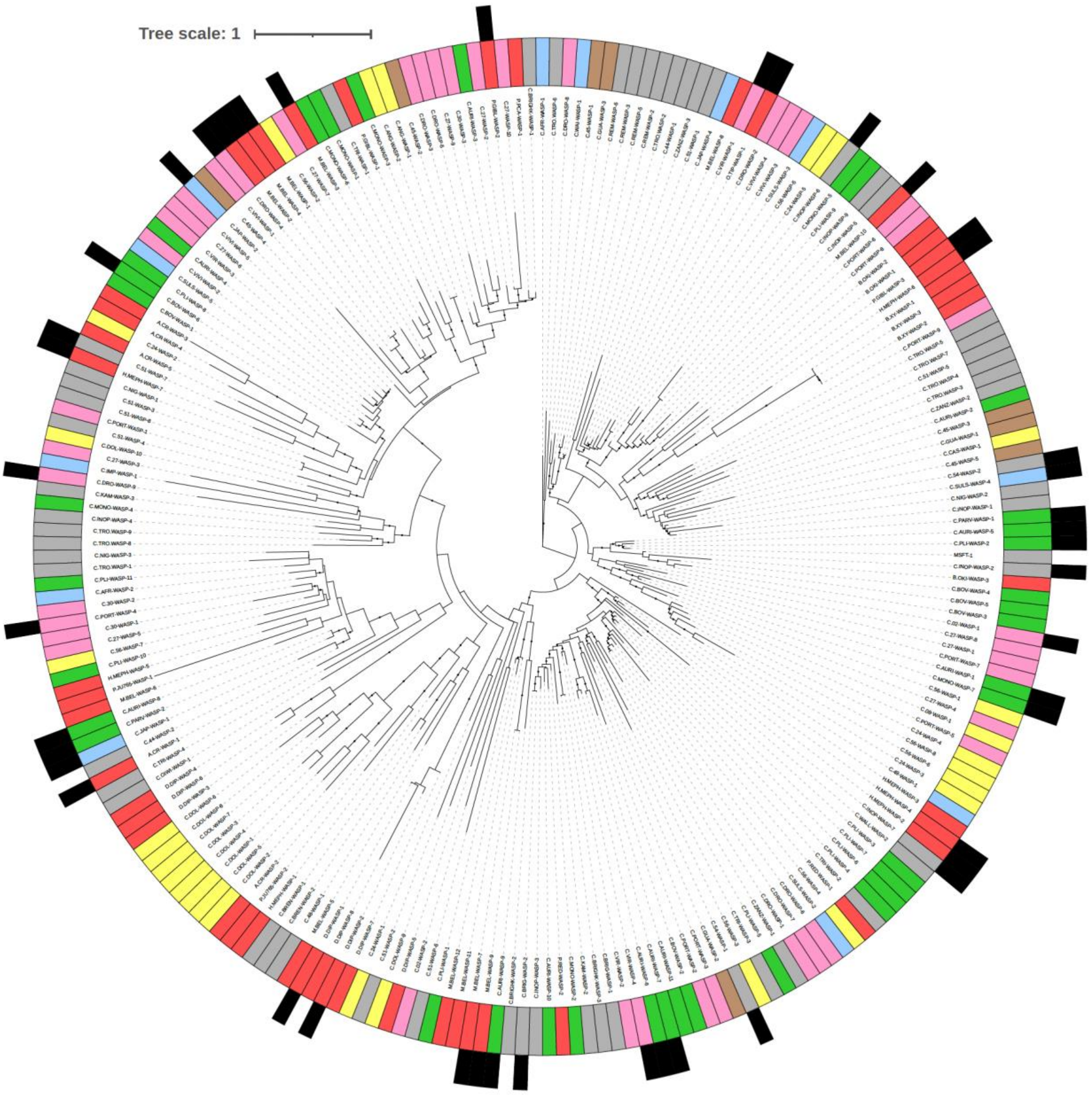


**Figure S10. Phylogenetic tree of WASP proteases.** Protein phylogenetic tree of 235 WASP proteases found across 52 nematode species. The *Caenorhabditis* group to which each ortholog belongs is represented by the box color of the inner ring (see Fig. 5A for color explanation). These groups are monophyletic except for the basal *Caenorhabditis* group (§) and the group containing non-*Caenorhabditis* species. Black boxes in the outer ring represent those WASP proteases that are associated with *Mavericks* in the genome. Black dots denote branches with a bootstrap value >85.

```
MSFT-1          1 MAQEIRIKMMGKDIAFELTDNRITKNSIKAGFLLEDDAVISLSYEKNAHQ  50
Cpli-WASP-2     1 MAQEIRIKMMDKDIAFELTDNRITKNSIKAGFLLEDDAVISLSYEKNAHQ  50

MSFT-1         51 ICCQMNEAGTSFLLPDEWSSMQFSVESDKAPSRPATPADIEGPNPKKRKL 100
Cpli-WASP-2    51 ICCQMNEAGTSFLLPDEWSSMQFSVESDKAPSRPATPADIEGPNPKKRKL 100

MSFT-1        101 DPYPEQLAVNGDLAASIGRYALYYNVKRDNYKRCAIPLTPRLAATFHHGE 150
Cpli-WASP-2   101 DPYSEQLAVNGDLAASIGRYALYYNVKKDNYKRCAIPLTPRLAATFHHGD 150

MSFT-1        151 NKSIKIGDEIVIHSWLDKNLKVATRAVKIMEEYDTIILQADGEHLSDKDF 200
Cpli-WASP-2   151 NKSIKIGDEIIIHSWLDKNLKVATRAVKILEEYDTIILQTDGEDLSDKDF 200

MSFT-1        201 ITSSGYPQKGMKYLLMGFSIIHEKTSHLSLSSGIIATEVTRRLRYTGSSG 250
Cpli-WASP-2   201 ISNSRYPRKGMKYLLMGFSIIHEKTSHLSLSSGIIATEVTRRLRYTGSSG 250

MSFT-1        251 SFKGDSGGGCWSEDGHLMGMQIGVEKVPHTKDTNGRPASPASGGRCCIIA 300
Cpli-WASP-2   251 SFKGDSGGGCWSEDGHLMGMQIEVEKVPHTKDTNGRPASPATGGRCCIIA 300

MSFT-1        301 IRDILGHIQDLMPMPADSDADSWSE                          325
Cpli-WASP-2   301 IRDILANIQDLMPMPADSDADSWSE                          325
```

**Figure S11. Protein alignment of MSFT-1 and *C.pli*-WASP-2.** There are a total of sixteen amino acid differences between *C. briggsae* MSFT-1 and *C. plicata* WASP-2, a cargo protease found in a *Maverick* transposon. The E273G is the only change at a site highly conserved in WASP protease evolution.

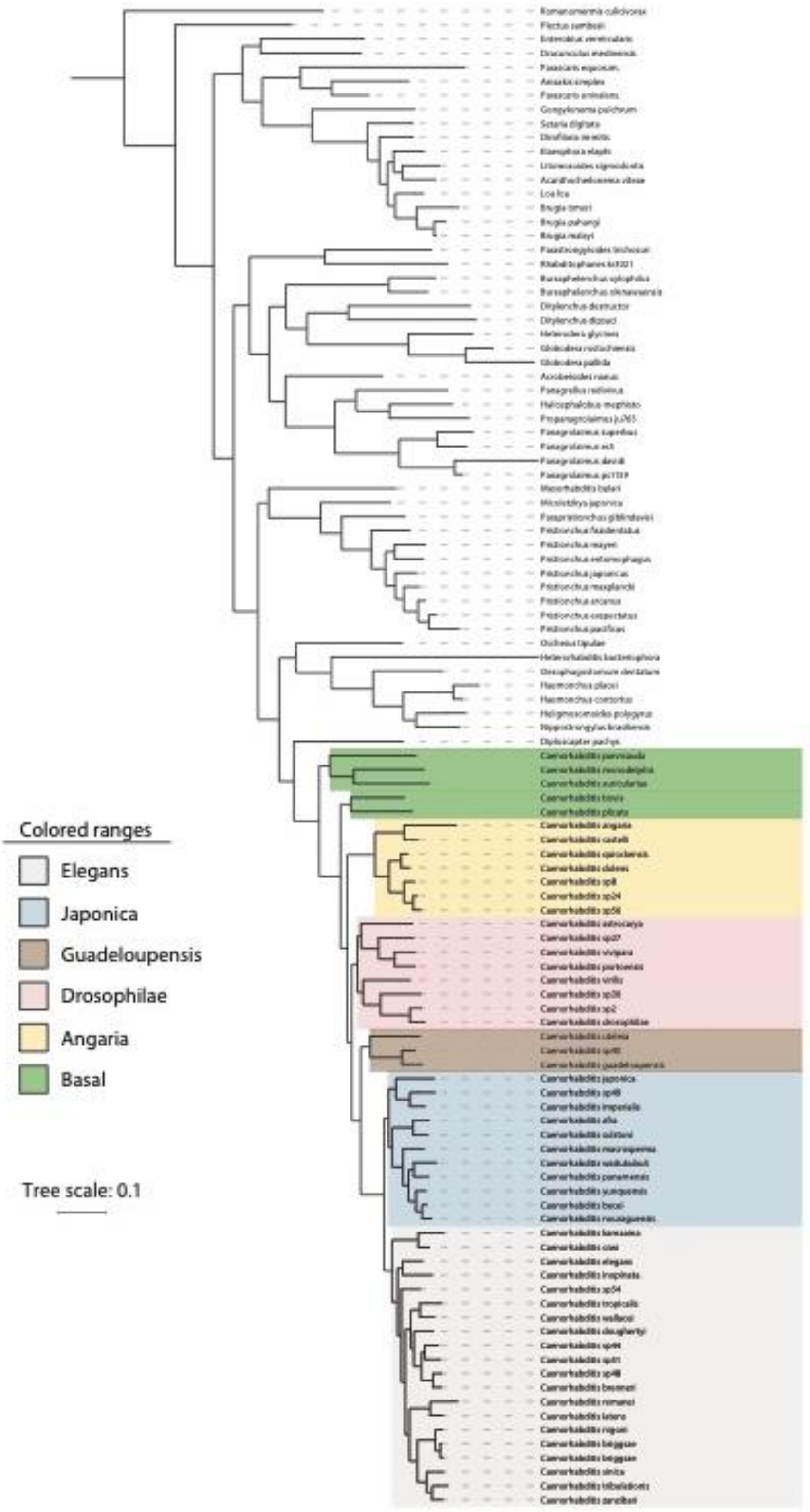


**Figure S12. Phylogenetic tree of nematode species used in this study.** The phylogenetic tree was obtained with Orthofinder. *Caenorhabditis* groups are highlighted with colors.

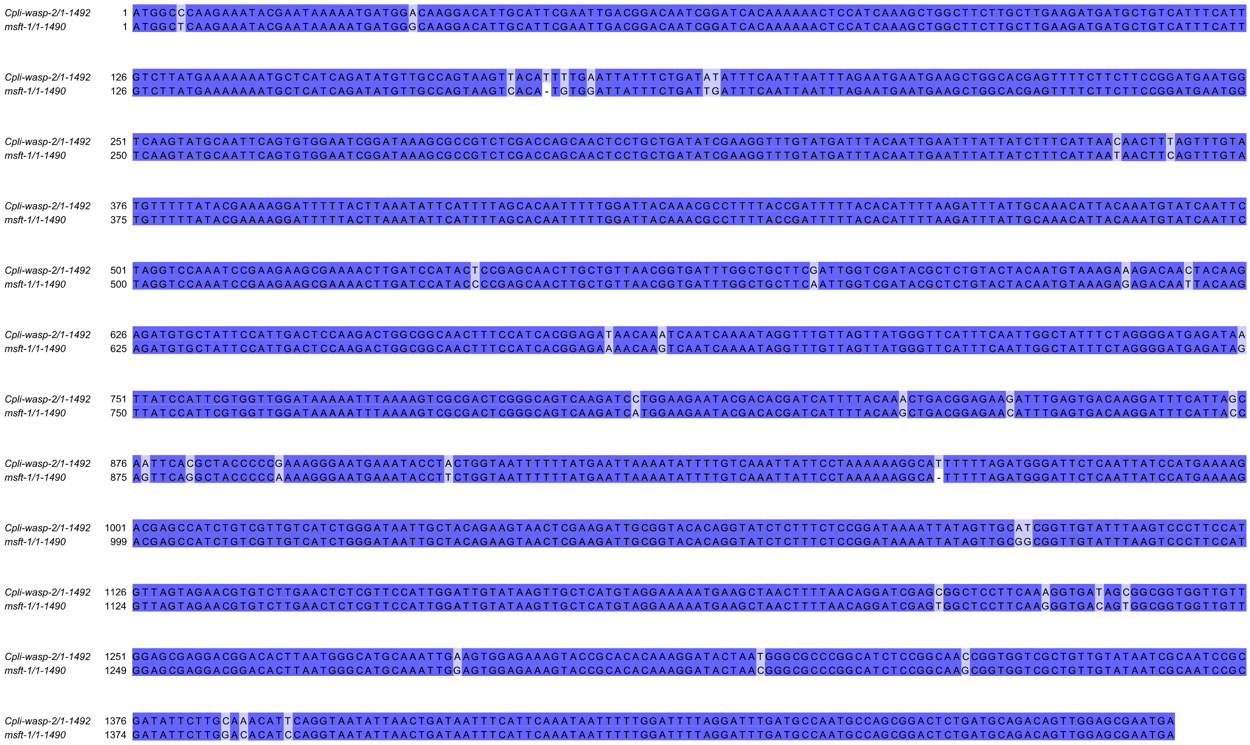

**Figure S13. Nucleotide alignment of *msft-1* and *C.pli-wasp-2*.** Aligned sequences include exons and introns.

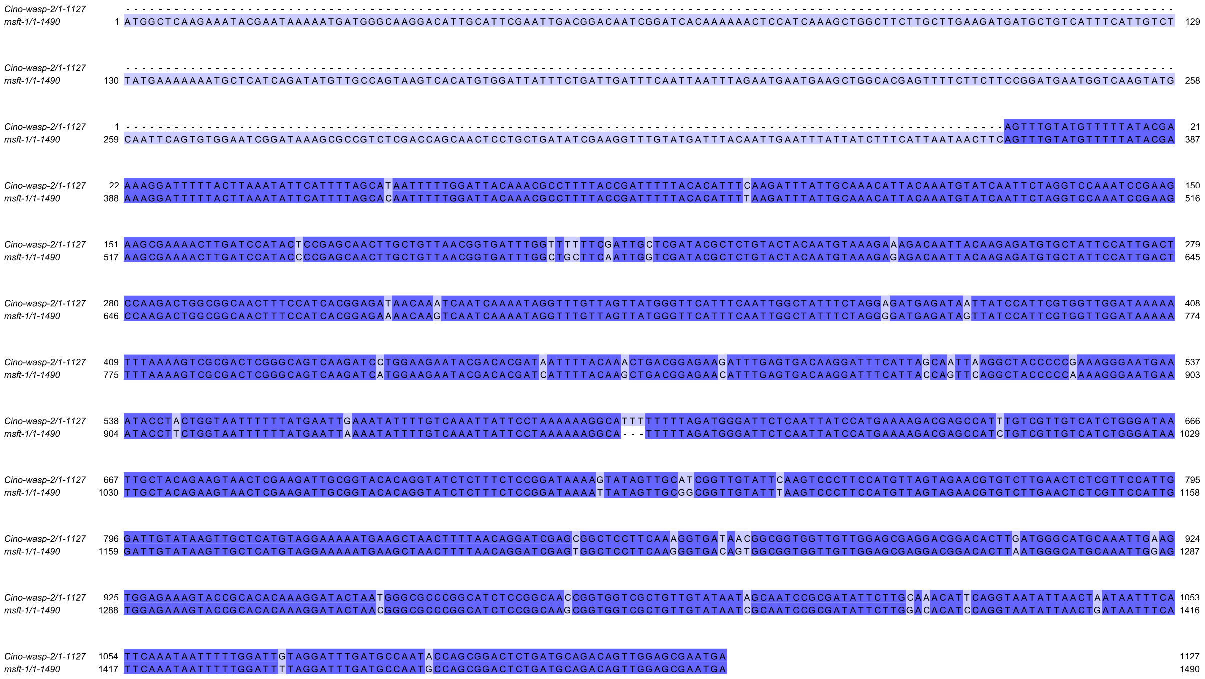


**Figure S14. Nucleotide alignment showing extreme conservation between *msft-1* and *Cino*-wasp-2.** Aligned sequences include exons and introns. *Cino*-wasp-2 has a premature stop codon, which is equivalent to a termination in MSFT-1 in amino acid position 203.

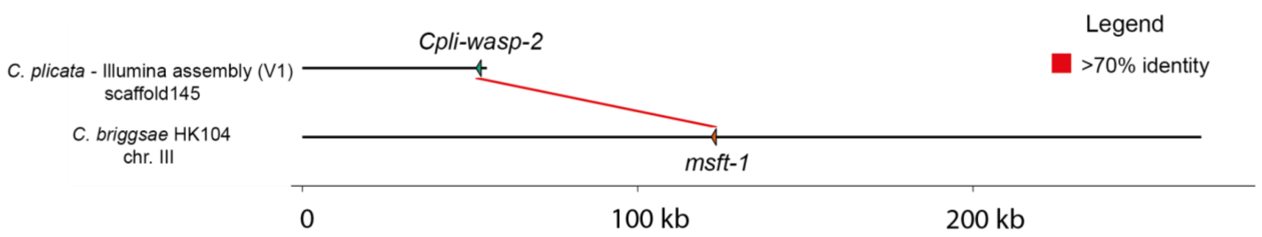


**Figure S15. Lack of homology between the neighboring regions of *msft-1* and *C.pli-wasp-2*.** Except for *C.pli-wasp-2* and *msft-1* there is no homology between the genomic regions containing these two genes in *C. plicata* and *C. briggsae* HK104.

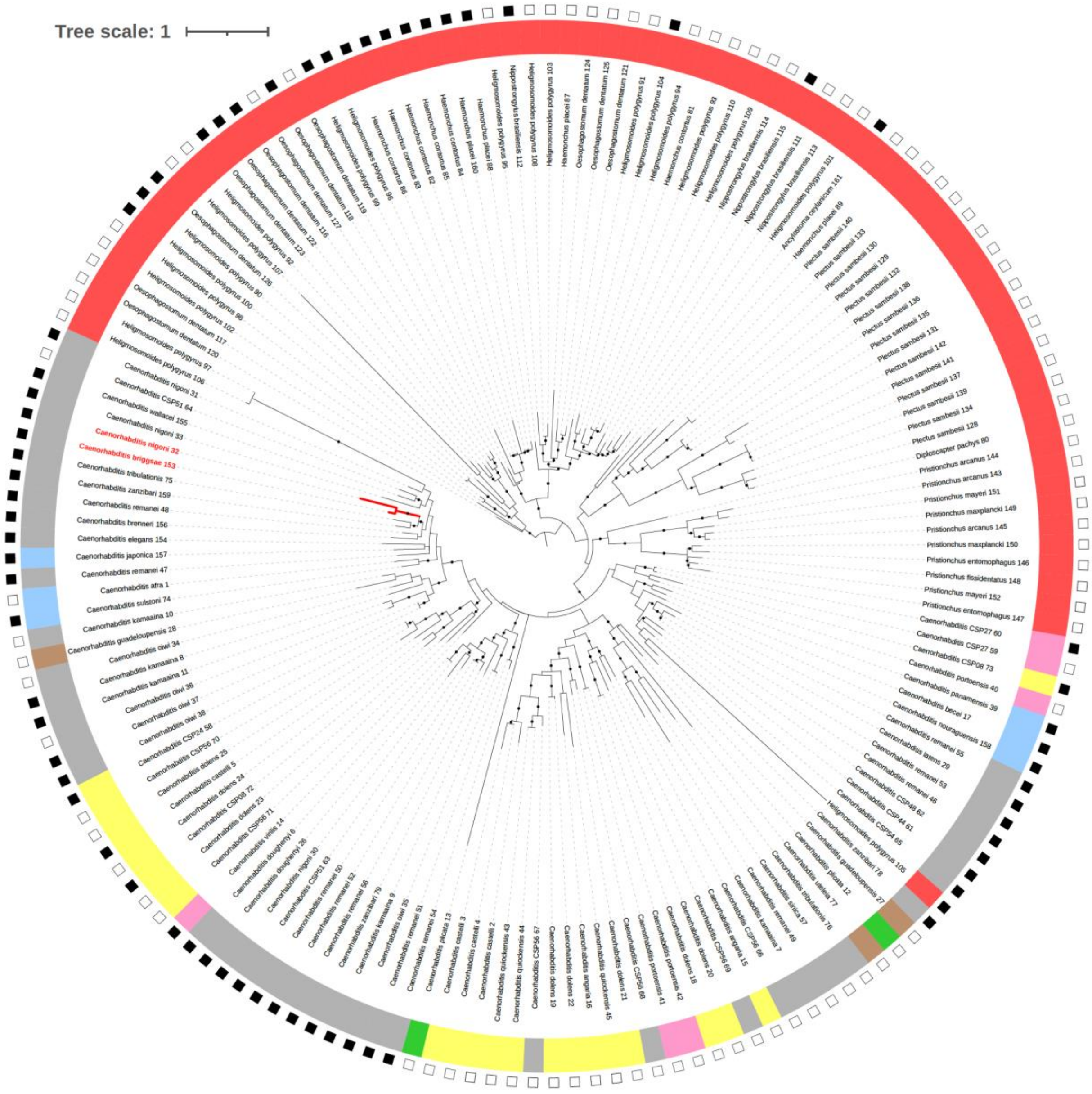


**Figure S16. Phylogenetic tree of the catalytic domain of the nematode MULE transposase.** Protein phylogenetic tree of 161 MULE-catalytic domains found across 52 nematode species. The *Caenorhabditis* group to which each ortholog belongs is represented by the box color of the inner ring (see Fig. 5A for color explanation). Non-*Caenorhabditis* species are represented by the red box color of the inner ring. Black outer squares represent those MULE transposases that are associated with *maap* (MULE-transposon associated proteins) genes in the genome. *C. briggsae* HK104 MULE transposase and its closest orthologue (*C. nigoni*) are highlighted in the phylogeny with red labels and red branches. Black dots denote branches with a bootstrap value >85. Midpoint rooting was used.

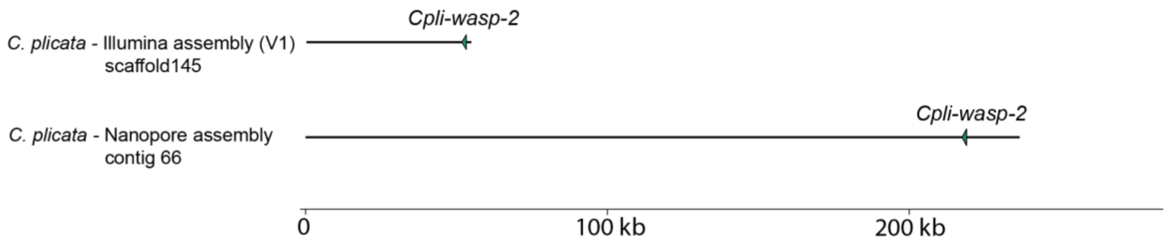


**Figure S17. Localization of *C.pli*-wasp-2 in fragmented *C. plicata* Illumina genome assembly, and *de novo* Nanopore-based *C. plicata* genome assembly.** In the *C. plicata* genome V1 (Illumina short-reads), the *Cpli-wasp-2* gene was found almost at the end of the Scaffold145, preventing us from the genomic neighborhood of *Cpli-wasp-2* in detail. This is likely not a coincidence but reflects a failure of the assembler in dealing with *Maverick* transposons that are in multiple copies in genomes. We observed a similar pattern across many low-quality genomes. To facilitate the study of *Cpli-wasp-2,* we generated a Nanopore long-read assembly of *C. plicata*. In this version of the genome, *Cpli-wasp-2* is not found at the edge of the scaffold, and we retrieved the full *Maverick*.

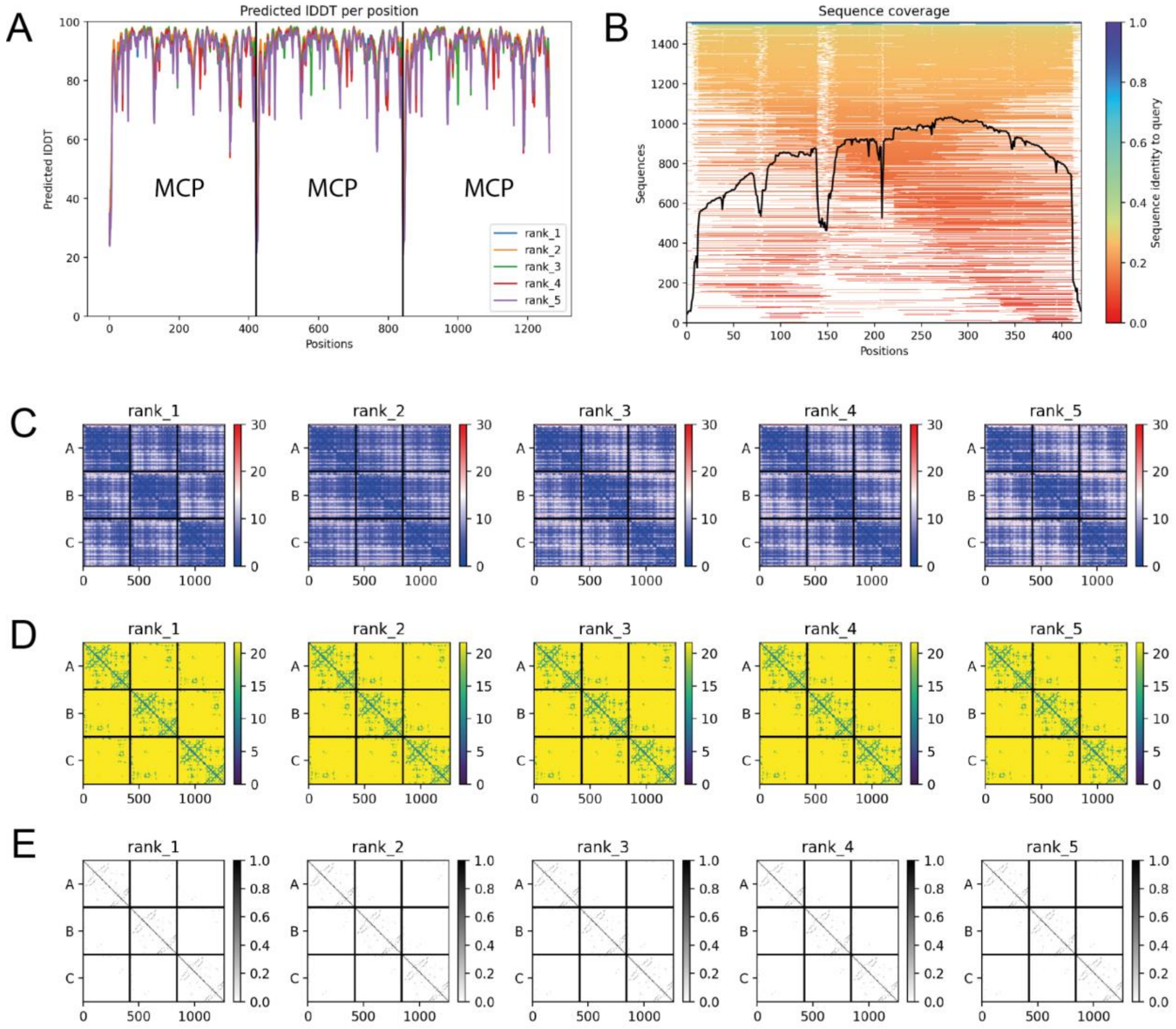


**Figure S18. Structural prediction of *Maverick* major capsid protein trimer. (A)** AlphaFold2 per-residue confidence metric pLDDT (Local Distance Difference Test) **(B)** Coverage of the multiple sequence alignment**. (C)** Predicted aligned error, confidence in the domain packing and large-scale topology of the protein. **(D)** Predicted distogram. **(E)** Predicted contacts.

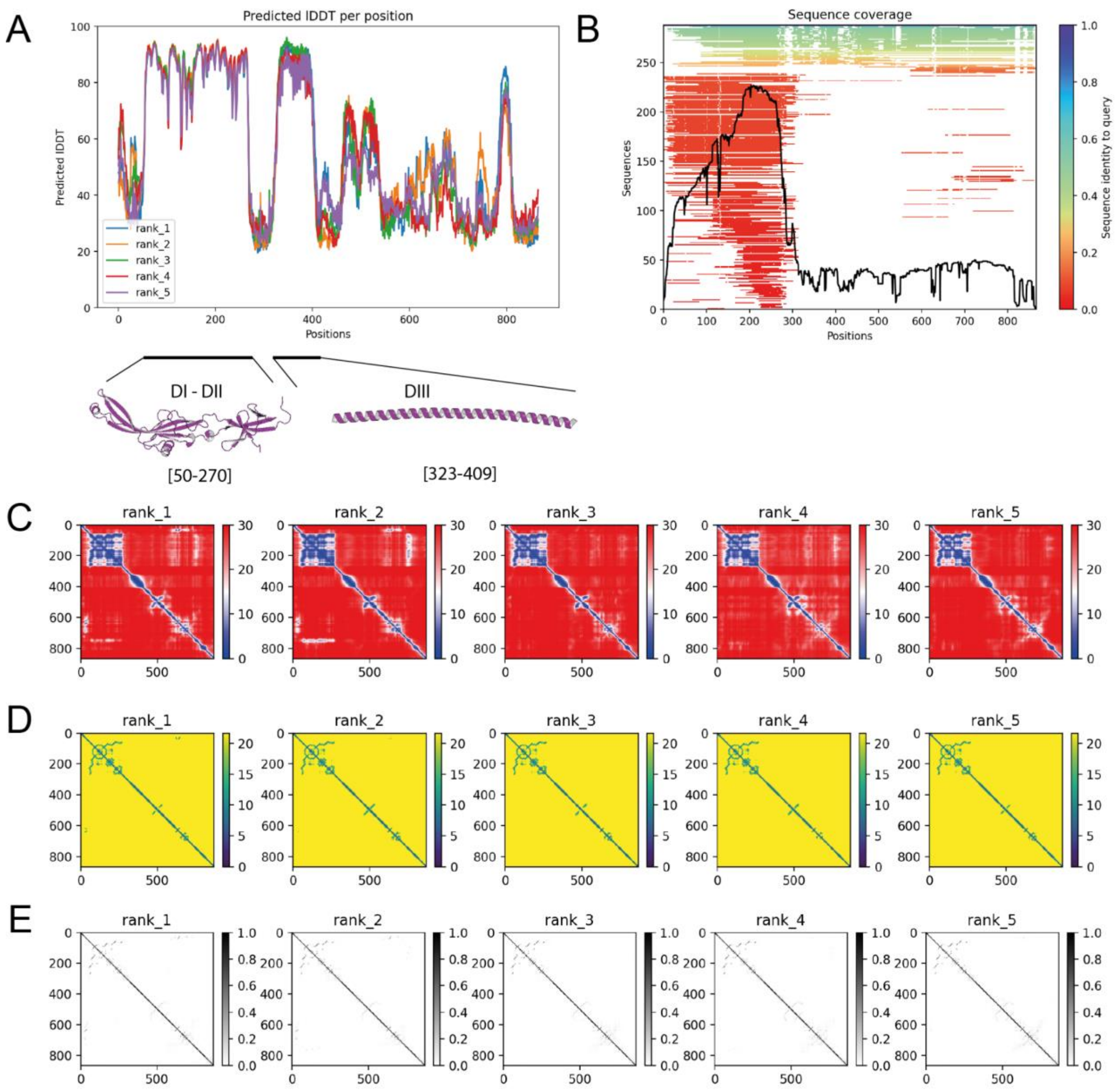


**Figure S19. Structural prediction of *Maverick* MFUS-1 fusogen. (A)** AlphaFold2 per-residue confidence metric pLDDT (Local Distance Difference Test). Only two large domains of MFUS-1 were predicted with high accuracy. A first domain from 50-270 AA, which includes the DI-DII of the fusogen and a region from 323-409 AA that likely represents DIII, a large α-helix that forms a trimeric coiled coil in viruses. **(B)** Coverage of the multiple sequence alignment. **(C)** Predicted aligned error, confidence in the domain packing and large-scale topology of the protein. **(D)** Predicted distogram. **(E)** Predicted contacts.

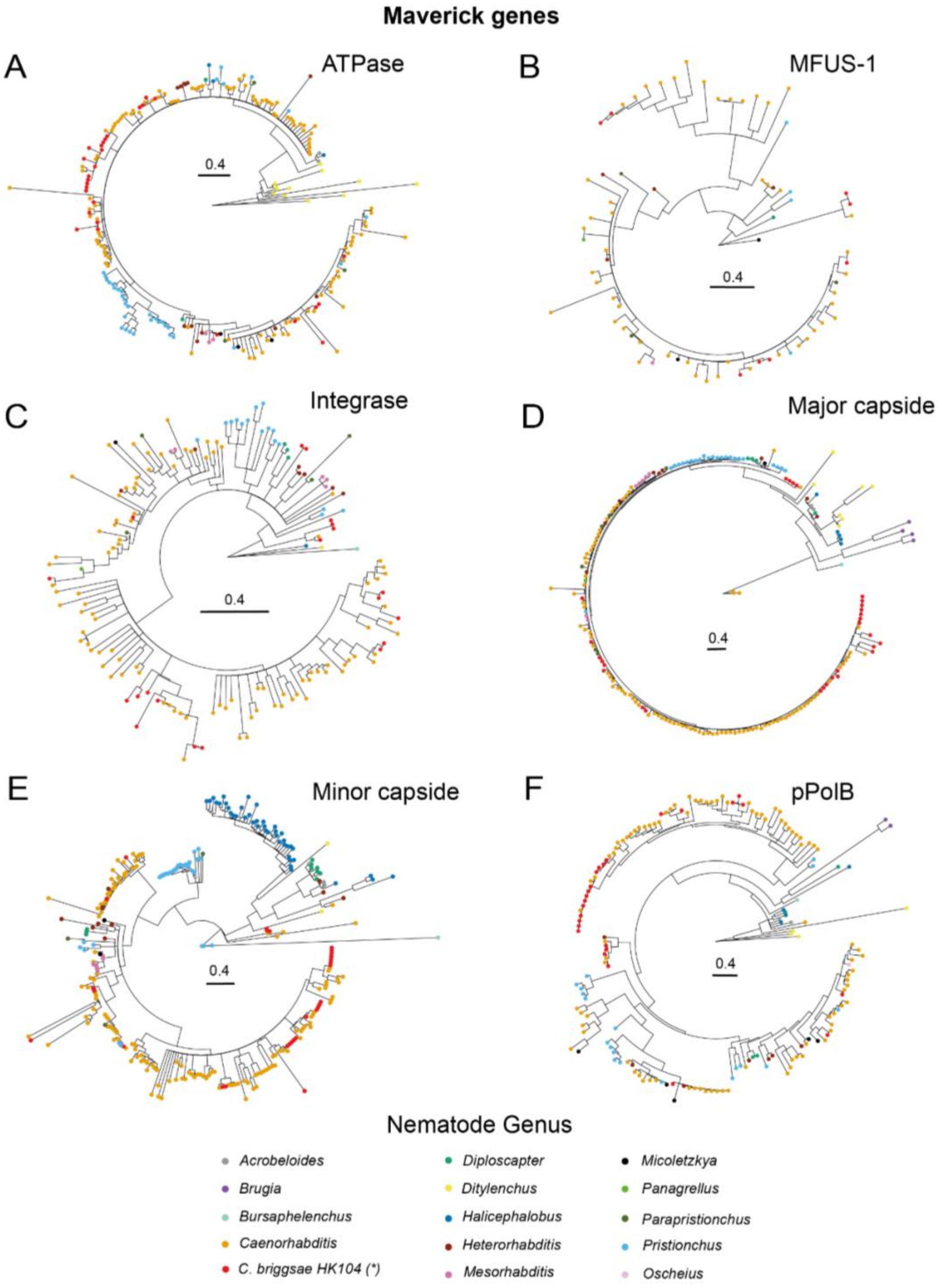


**Figure S20. Phylogenetic protein tree of nematode *Maverick* genes.** Maverick genes: **(A)** ATPase. **(B)** MFUS-1 (fusogen). **(C)** Integrase. **(D)** Major capside. **(E)** Minor capside. **(F)** pPolB.Terminal nodes color represents nematode genus. Branches with boostrap < 75 were collapsed. We filtered out those sequences that were shorter (<60%) than the consensus Maverick gene except for pPolB (sequences below 500 bp length were filtered out).

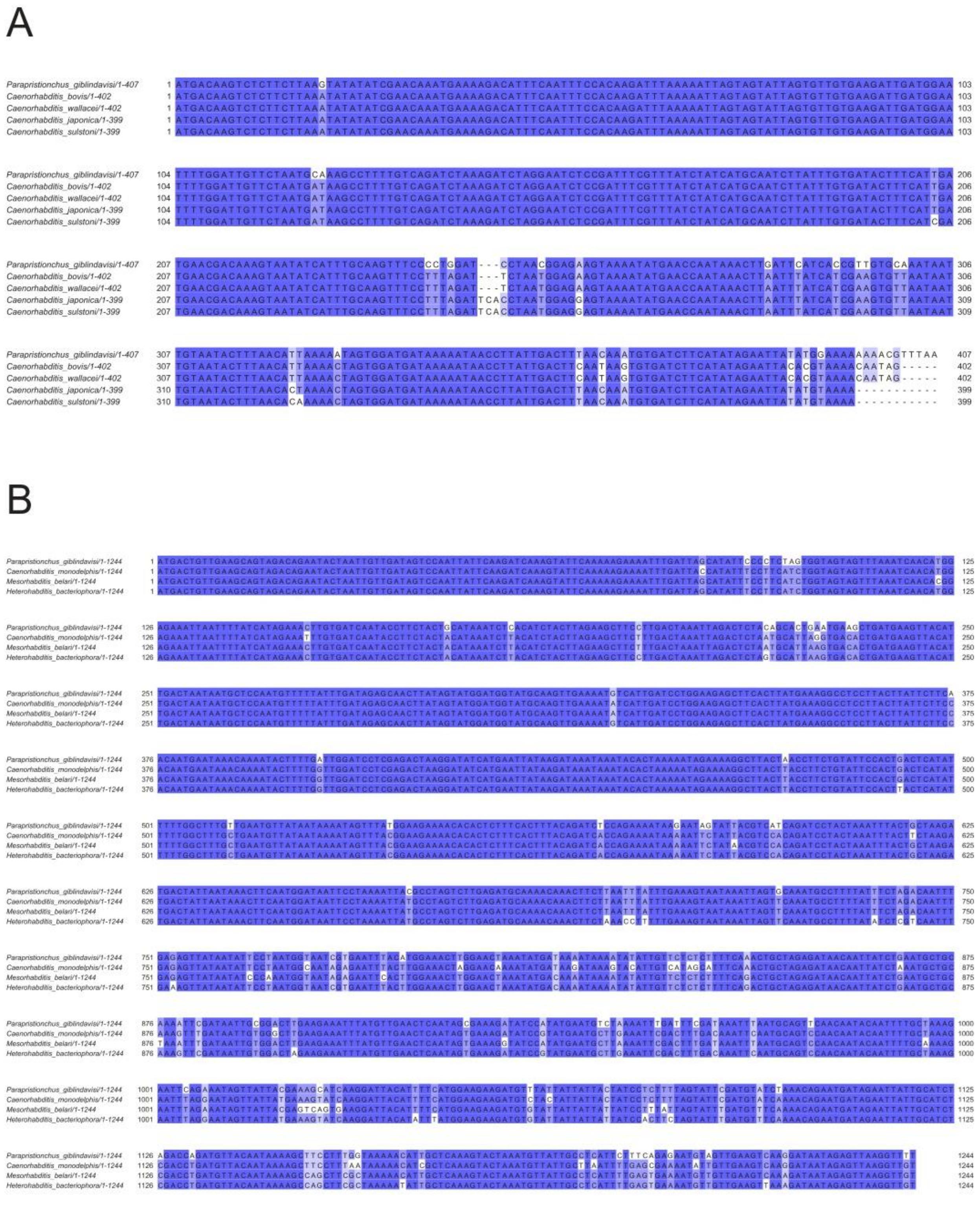

**Figure S21. Nucleotide alignment of selected *Maverick* gene. (A)** *Maverick* major capsid gene **(B)** *Maverick* minor capsid gene. In both cases the nucleotide identity is ~90-95% between different species which belong to different genus: *Caenorhabditis*, *Heterorhabditis*, *Mesorhabditis*, *Parapristionchus*.

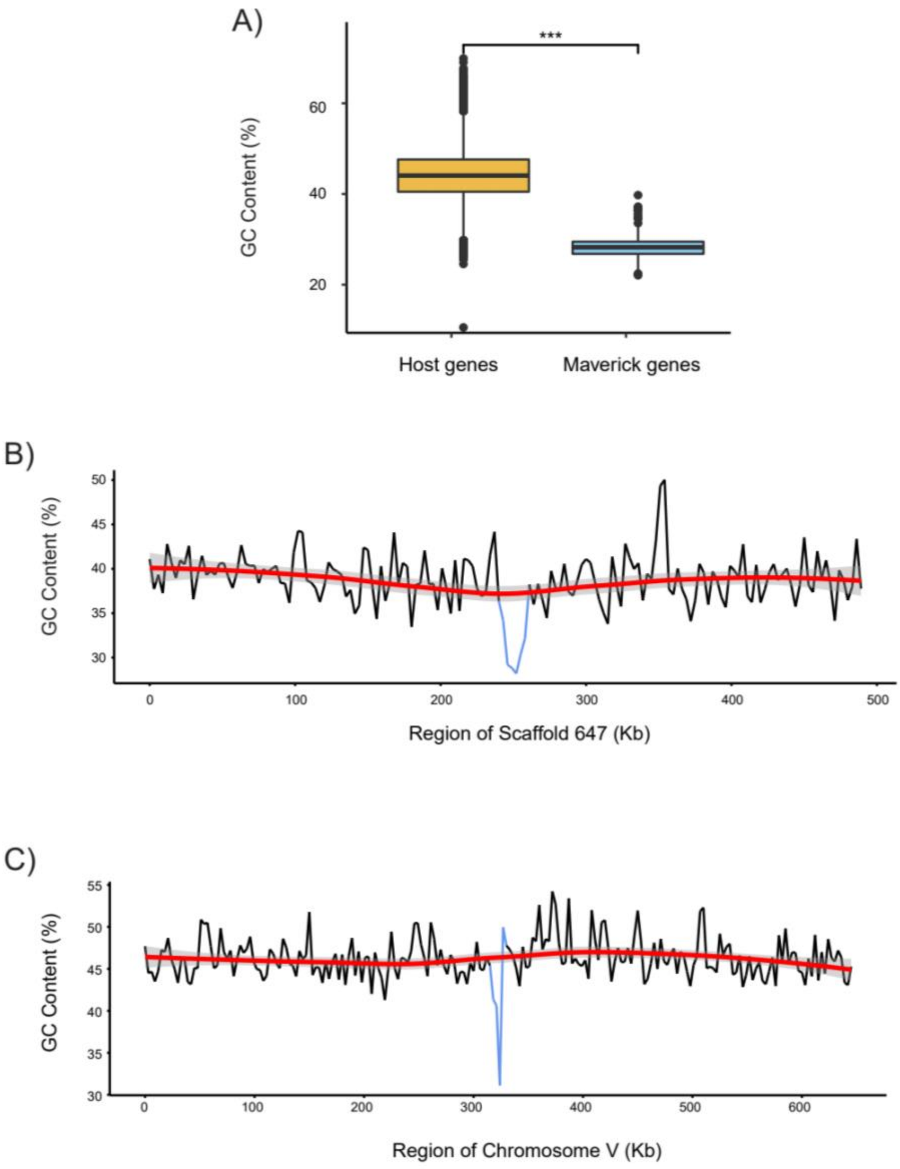


**Figure S22. Differences in GC content between nematode and *Maverick* genomes. (A)** Differences in GC content between *Maverick* genes (n=243) and nematode core-genes (n=74,655). Species used in this analysis are *C. briggsae* HK104, *C. plicata*, *C. parvicauda* and *C. auriculariae.* **(B)** GC content of *C. auriculariae* scaffold 647 region (sliding window=3kb) **(C)** GC content of *O. tipulae* Chr. V region (sliding window=3kb). A *Maverick* copy is located in the blue region in both B and C.

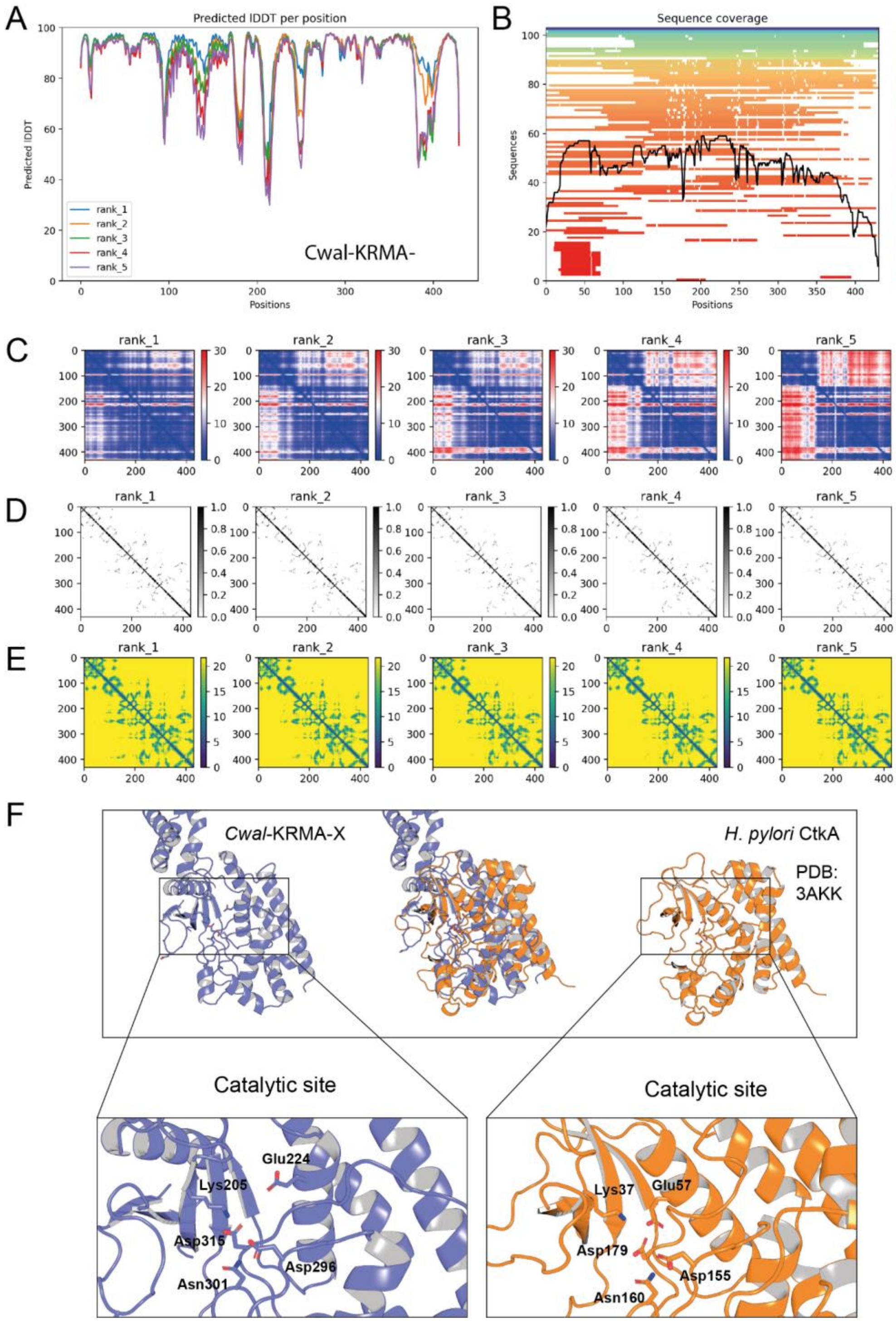


**Figure S23. Structural prediction of the *Cwal*-KRMA-1 kinase. (A)** Alphafold2 per-residue confidence metric pLDDT (Local Distance Difference Test). **(B)** Coverage of the multiple sequence alignment. **(C)** Predicted aligned error. **(D)** Predicted distogram. **(E)** Predicted contacts. **(F)** Comparison of *Cwal*-KRMA-1 model and a *H. pylori* CtkA structure generated by X-ray diffraction (PDB: 3AKK). Highlighted are those residues that make up the kinase catalytic site of CtkA and the inferred catalytic residues of *Cwal*-KRMA-1 based on structural homology.

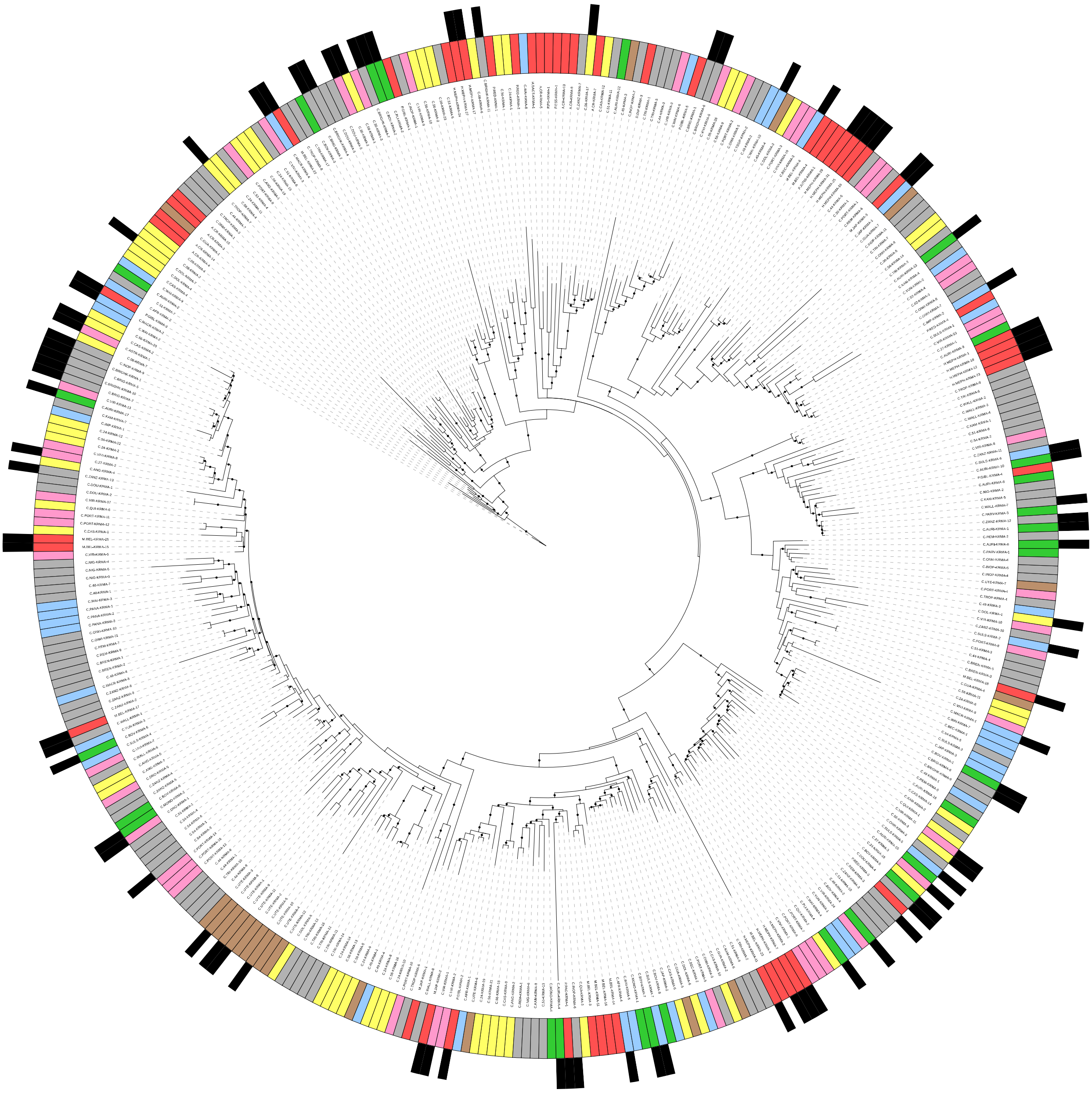

**Figure S24. Phylogenetic tree of nematode KRMA kinases.** The total number of KRMA kinases found in the KRMA Orthogroup was 532. However, given the high divergence of some kinases we used 370 KRMA kinases (found across 61 nematode species) for the phylogenetic reconstruction. The *Caenorhabditis* group to which each ortholog belongs is represented by the color of the inner ring. These groups are monophyletic except for the basal *Caenorhabditis* group and the group containing *non-Caenorhabditis* species. Black outer ring represents those KRMA kinases that are associated with Mavericks in the genome. Black dots denote branches with a bootstrap value >85.

**Table S1**. $F_2$ genotypes of relevant *C. briggsae* crosses. For an explanation of which crosses were genotyped, see Methods. AF=AF16, HK=HK104, n.g.=not genotyped.

| Figure # | | Cross | | | | Wildtype | | | | | Delayed | | | | | Embryonic/larval arrest | | | | | |
|---|---|---|---|---|---|---|---|---|---|---|---|---|---|---|---|---|---|---|---|---|---|
| | Description | Hermaphrodite | Male | Type | Total F2s | Total | n.g. | AF/AF | Het | HK/HK | Total | n.g. | AF/AF | Het | HK/HK | Total | n.g. | AF/AF | Het | HK/HK | Genotyped with (see Table S5) |
| S1 | Test cross with NIL1 for presence of TA element within the introgression | INK199 | AF16 | F1 self | 281 | 230 | 85 | 13 | 79 | 53 | 50 | 0 | **50** | 0 | 0 | 1 | 0 | 0 | 0 | 1 | Marker 2; AFLP |
| S1 | Test cross with NIL2 for presence of TA element within the introgression | INK200 | AF16 | F1 self | 79 | 68 | 0 | 8 | 39 | 21 | 11 | 0 | **11** | 0 | 0 | 0 | 0 | 0 | 0 | 0 | Marker 2; AFLP |
| S1 | Test if TA element is within common introgressed region in NIL1 and NIL2 | INK200 | INK199 | F1 self | 150 | 142 | 71 | 18 | 28 | 25 | 3 | 3 | **0** | 0 | 0 | 5 | 2 | 0 | 1 | 2 | Marker 2; AFLP |
| S1 | NIL1 maternal backcross to AF16 to confirm maternal effect of toxin | INK199 | AF16 | Maternal BC | 123 | 61 | 0 | 10 | 51 | N/A | 48 | 0 | **48** | 0 | N/A | 14 | 6 | 6 | 2 | N/A | Marker 2; AFLP |
| 6 | Cross to test effect of MSFT-1(K29A) mutation on toxicity/F2 delay | INK502 | AF16 | F1 self | 254 | 184 | 0 | 4 | 117 | 63 | 5 | 0 | 1 | 4 | 0 | 65 | 35 | **23** | 6 | 1 | Marker 2; AFLP |
| | | | | | | Total | n.g. | Mutant | Het | WT | Total | n.g. | Mutant | Het | WT | Total | n.g. | Mutant | Het | WT | |
| 1 | Cross with *msft-1 tlpr-1* double mutants to test for susceptibility to wildtype TA element | INK361 | INK199 | F1 self | 158 | 113 | 3 | 4 | 66 | 40 | 36 | 1 | **34** | 1 | 0 | 9 | 6 | 0 | 1 | 2 | *tlpr-1;* Sanger sequencing |
| text | Cross with *msft-1 tlpr-1* double mutants to test for susceptibility to wildtype TA element (second, independent antidote allele) | INK199 | INK480 | F1 self | 80 | 57 | 0 | 3 | 36 | 18 | 19 | 0 | **19** | 0 | 0 | 4 | 3 | 0 | 0 | 1 | *tlpr-1;* AFLP |

**Table S2**. List of genes within the candidate region containing the HK104 TA element.

| ORF number | Absent/divergent in AF16 | Expression high in HK104, low in AF16 | Is in tight linkage to another candidate gene | Scaffold | Start (bp) | End (bp) |
|---|---|---|---|---|---|---|
| 14760 | no | no | no | 4 | 5128510 | 5132903 |
| 14761 | no | no | no | 4 | 5133695 | 5137284 |
| 14762 | no | no | no | 4 | 5137530 | 5140494 |
| 14763 | no | no | no | 4 | 5140753 | 5141833 |
| 14764 | no | no | no | 4 | 5141980 | 5144526 |
| 14765 | no | no | no | 4 | 5144927 | 5148422 |
| 14766 | no | no | no | 4 | 5148988 | 5155142 |
| 14767 | **absent** | **yes** | **yes** | 4 | 5156573 | 5158038 |
| 14768 | **absent** | **yes** | **yes** | 4 | 5158761 | 5159174 |
| 14769 | no | no | no | 4 | 5164712 | 5165643 |
| 14770 | no | no | no | 4 | 5165699 | 5168349 |
| 14771 | no | no | no | 4 | 5169046 | 5170835 |
| 14772 | no | no | no | 4 | 5171233 | 5174550 |
| 14773 | no | no | no | 4 | 5174856 | 5177876 |
| 14774 | no | no | no | 4 | 5180706 | 5181275 |
| 14775 | no | no | no | 4 | 5181455 | 5187184 |
| 14776 | no | no | no | 4 | 5189257 | 5190133 |
| 14777 | no | no | no | 4 | 5192597 | 5193900 |
| 14778 | no | no | no | 4 | 5194584 | 5195803 |
| 14779 | no | no | no | 4 | 5198563 | 5210403 |
| 14780 | no | no | no | 4 | 5210758 | 5212365 |
| 14781 | no | no | no | 4 | 5212463 | 5214367 |
| 14782 | no | no | no | 4 | 5213971 | 5214319 |
| 14783 | no | no | no | 4 | 5214417 | 5215199 |
| 14784 | no | no | no | 4 | 5215550 | 5218096 |
| 14785 | no | no | no | 4 | 5219965 | 5220135 |
| 14786 | no | no | no | 4 | 5225080 | 5236192 |
| 14787 | no | no | no | 4 | 5238597 | 5239317 |
| 14788 | no | no | no | 4 | 5255809 | 5256683 |
| 14789 | no | no | no | 4 | 5258871 | 5261045 |
| 14790 | no | no | no | 4 | 5261063 | 5262071 |
| 14791 | no | no | no | 4 | 5262337 | 5263564 |
| 14792 | no | no | no | 4 | 5263655 | 5264474 |
| 14793 | no | no | no | 4 | 5264841 | 5266541 |
| 14794 | no | no | no | 4 | 5268755 | 5270776 |
| 14795 | no | no | no | 4 | 5270981 | 5273168 |
| 14796 | no | no | no | 4 | 5274661 | 5278168 |
| 14797 | no | no | no | 4 | 5285740 | 5286528 |
| 14798 | no | no | no | 4 | 5289084 | 5291135 |
| 14799 | no | no | no | 4 | 5291730 | 5293365 |
| 14800 | no | no | no | 4 | 5293451 | 5294345 |
| 14801 | no | no | no | 4 | 5294996 | 5296176 |
| 14802 | no | no | no | 4 | 5298454 | 5301778 |
| 14803 | no | no | no | 4 | 5302165 | 5302577 |
| 14804 | no | no | no | 4 | 5302745 | 5303668 |

**Table S3**. Nematode strains used in this study.

| Strain name | Species | Genotype | Description | Source |
|---|---|---|---|---|
| AF16 | *C. briggsae* | Wildtype (reference) | Wild isolate from Ahmedabad, Gujarat, India. Collected by A. Fodor. | CGC |
| HK104 | *C. briggsae* | Wildtype | Wild isolate from Okayama, Japan. Collected by S. Baird. | CGC |
| JU439 | *C. briggsae* | Wildtype | Wild isolate from Reykjavík, Iceland. Collected by A. Barrière. | CGC |
| QR24 | *C. briggsae* | Wildtype | Wild isolate from Montreal, Quebec, Canada. Collected by C. Rocheleau. | CGC |
| ED3036 | *C. briggsae* | Wildtype | Wild isolate from Tien Mu, Taipei, Taiwan. Collected by A. Cutter. | CGC |
| INK199 | *C. briggsae* | *abuIR5*(III:2.98-7.90 Mb, HK104 > AF16) | NIL 1, derived from 16x maternal backcross to AF16 | This study |
| INK200 | *C. briggsae* | *abuIR6*(III:4.0-9.8 Mb, HK104 > AF16) | NIL 2, derived from 16x maternal backcross to AF16 | This study |
| INK202 | *C. briggsae* | | 16x HK->AF16 maternal backcross | This study |
| INK302 | *C. briggsae* | *msft-1(abu315*[MSFT-1[S77X]+MSFT-1[S77X]]) III; *abuIR5* | *msft-1* HDR (c.219CAGTGTGGAATCG>AAGCTTTGAATΔ[CG]). Gene was also duplicated during creation of the mutant: second locus is linked but location unknown, genotype is HDR+122bp insertion. Resulting proteins identical, p.S77X. INK199 background. | This study |
| INK361 | *C. briggsae* | *msft-1(abu315*[MSFT-1[S77X]+MSFT-1[S77X]]) III; *tlpr-1*(*abu317*[TLPR-1[E87GfsX2]]) III; *abuIR5* | *tlpr-1* 2bp deletion (c.258GGAA>Δ[GG]AG) = 2 residue substitution, premature stop (p.EG87-88GV, F89X). In INK302 (MSFT-1 S77X), INK199 background. | This study |
| INK390 | *C. briggsae* | *msft-1(abu319*[MSFT-1[S256A]]) III; *abuIR5* | *msft-1* (c.AG766-767GC), p.S256A | This study |
| INK391 | *C. briggsae* | *msft-1(abu320*[MSFT-1[G273E]]) III; *abuIR5* | *msft-1* (c.G817A, c.G822T), p.G273E | This study |
| INK465 | *C. briggsae* | *msft-1(abu319*[MSFT-1[S256A]]) III; *abuIR5* | *msft-1* (c.AG766-767GC), p.S256A. Independently derived from INK390. | This study |
| INK467 | *C. briggsae* | *msft-1(abu320*[MSFT-1[G273E]]) III; *abuIR5* | *msft-1* (c.G817A, c.G822T), p.G273E. Indepenently derived from INK391. | This study |
| INK480 | *C. briggsae* | *msft-1(abu315*[MSFT-1[S77X]+MSFT-1[S77X]]) III; *tlpr-1(abu318*[TLPR-1[K65X]]) III; *abuIR5* | *tlpr-1* HDR2 (c.A193T, c.Δ197-245, c.C246T, c.AT250-251TA). Premature stop (p.K65X). | This study |
| INK502 | *C. briggsae* | *msft-1(abu321*[MSFT-1[K29A]]) III; HK104 | *msft-1* (c.CAA84-86TGC), p.K29A. HK104 background. | This study |
| SB355 | *C. plicata* | Wildtype | Wild isolate from Berlin, Germany. Collected by J. Völk. | Christian Braendle |

**Table S4**. sgRNAs and HDR templates used to generate mutant worm lines in this study.

| Gene | Purpose | Guide 1 | Guide 2 | Repair template sequence | Lines generated |
|---|---|---|---|---|---|
| *msft-1* | Premature stop | TATGCAATTCAGTGTGGAAT | - | caattaatttagAATGAATGAAGCTGGCACGAGTTTTCTTCTTCCGGATGAATGGTCAAGTATGCAATTaAGcttTGAATGATAAAGCGCCGTCTCGACCAGCAACTCCTGCTGATATCGAAGgtttgtatgatttacaattg | INK302 |
| *msft-1* | S256A | TCCTTCAAGGGTGACAGTGG | - | CTCCACTCCAATTTGCATGCCCATTAAGTGTCCGTCCTCGCTCCAACAACCACCGCCAgcGTCACCCTTGAAGGAGCCACTCGATCctgttaaaagttagcttcatttttcctacatgagc | INK390, INK465 |
| *msft-1* | G273E | AATGGGCATGCAAATTGGAG | - | GCGACCACCGCTTGCCGGAGATGCCGGGCGCCCGTTAGTATCCTTTGTGTGCGGTACTTTCTCaACTtCAATTTGCATGCCCATTAAGTGTCCGTCCTCGCTCCAACAACCACCGCCACTGTCACCCTTGAAGGAGCC | INK391, INK467 |
| *msft-1* | K29A | AAGCAAGAAGCCAGCTTTGA | - | GGCAAGGACATTGCATTCGAATTGACGGACAATCGGATCACAAAAAACTCCATtgcAGCTGGCTTCTTGCTTGAAGATGATGCTGTCATTTCATTGTCTTATGAAAAAAATGC | INK502 |
| *tlpr-1* | Premature stop | ATAGATGGGTCTTTATGGAC | ACCCCGACATGCACGAGGAA | CACAGGAAGTGTACCGGTTGGAGGCATTCCTTGCATCTCCGCCATTTTGTGAACTGGGCATCAAACCCCTTagtcacttaGACTGGAATGACACTAGAAGAACGTGCTTCCTTACGCTGCTTTTTTCTTCTctaaaattgttcaatttag | INK361 |
| *tlpr-1* | Premature stop | CATACATTGCAAATAGATGGG | AGAGGGACCCAGAGCTCGACC | CCGGTTGGAGGCATTCCTTGCATCTCCGCCATTtTGTGAACTGGGcATCAAaCCCTTCCTCGTGCtagtcacttaATGGACTGGAATGAcACTAGAAGAACGTGCTTCCTTACGCTGCTTTTTTCTTCTctaaaattgttc | INK480 |

**Table S5**. Genotyping primers and assays used in this study.

| Marker or gene | Forward primer | Reverse primer | Position (AF16) | Amplicon (bp) | Position (HK104) | Amplicon (bp) | Assay | Comments |
|---|---|---|---|---|---|---|---|---|
| _Genotyping of mutations and crosses:_ | | | | | | | | |
| Marker 2 | GGCGATTTTGTTAGTTCAAA | AATTCAGGTTCTTACAGGGATAC | III:4,012,867-4,013,114 | 249 | 4:4,148,047-4,148,321 | 293 | PCR, AFLP | Used to genotype all crosses between AF16 and NIL background lines. |
| *msft-1* (A) | TCGCTTCTTCGGATTTGGACCT | CGTCGCTCAAAATGGCTCAAGA | N/A | N/A | 4:5,157,516-5,158,049 | 533 | PCR, Sanger seq | *msft-1(ko)*, K29A genotyping (R primer to sequence) |
| *msft-1* (B) | TTTCATTCGCTCCAACTGTCTGC | GGATCTGTCGTTGTCATCTGGGA | N/A | N/A | 4:5,156,571-5,157,058 | 487 | PCR, Sanger seq | *msft-1* S256A, G273E genotyping (R primer to sequence) |
| *tlpr-1* | GCCTGCTACTATAGATTTTCATGCCC | ACGTGTTTCAGGACCTCGATTCC | N/A | N/A | 4:5,158,371-5,159,410 | 1039 | PCR, AFLP/seq | *tlpr-1(ko)* genotyping (R to sequence) |
| _Chr. III genotyping markers across introgression (Fig. S1):_ | | | | | | | | |
| Marker 1 | TAAGGTAGGTGGCGTTTAAAGGCG | GAATTTCGACCAGTGCGTTGCGG | III:2,988,036-2,988,128 | 468 | 4:2,957,338-2,957,713 | 374 | PCR, AFLP | This study |
| Marker 2 | GGCGATTTTGTTAGTTCAAA | AATTCAGGTTCTTACAGGGATAC | III:4,012,867-4,013,114 | 249 | 4:4,148,047-4,148,340 | 293 | PCR, AFLP | Indel ID: bhP38 (Koboldt et al. 2010) |
| Marker 3 | TTGCTGCAGGTTTATATCACTA | ACTTGAAGATCTGTTGACTGGT | III:5,752,834-5,753,060 | 248 | 4:5,824,955-5,825,173 | 219 | PCR, AFLP | Indel ID: bhP12 (Koboldt et al. 2010) |
| Marker 4 | TGTCTGCATTTTTGGAAAGTTCTGC | TAAATGCGATTTGGCTCTAACCGG | III:6,253,922-6,255,018 | 477 | 4:6,316,435-6,316,808 | 380 | PCR, AFLP | This study |
| Marker 5 | TAGTTTTTAACTGCTCCAGAACT | TGTATGGAAGTCTTTTGAAGAAA | III:7,747,668-7,747,945 | 300 | 4:7,884,674-7,884,990 | 283 | PCR, AFLP | Indel ID: cb-s272 (Koboldt et al. 2010) |
| Marker 6 | TCCTCGACAGTTGGTCCTTGTGGG | CACCGTTCTACCGCATTTTTCTCGAC | III:7,933,218-7,933,321 | 292 | 4:8,101,757-8,101,941 | 187 | PCR, AFLP | This study |
| Marker 7 | GGGGGCTGTCTGATTACTGTAACG | GCTTAGACGCGAATATTTTCAATCTCG | III:8,823,010-8,823,053 | 245 | 4:8,996,236-8,996,435 | 200 | PCR, AFLP | This study |
| Marker 8 | CTTGGTATGCCAACATTTTTAT | AAAATAGTCAGATTCTCAGTCCA | III:9,844,512-9,845,084 | 573 | 4:9,982,731-9,983,207 | 477 | PCR, AFLP | Indel ID: cb-m54 (Koboldt et al. 2010) |
| Marker 9 | AATCTAGAAGGTTTTTGGTTTTT | ACTGTCGTTCCTTGTATTTTTC | III:11,365,281-11,365,990 | 710 | 4:11,755,177-11,755,780 | 813 | PCR, AFLP | Indel ID: cb-m157 (Koboldt et al. 2010) |
| _Chr. III markers to narrow candidate region (Fig. 1C):_ | | | | | | | | |
| Marker 2b | AGAATTTCAAAAAGACGGCACAC | TTTTCCTTTTTCATGAAGAGTTTTG | III:4,888,337-4,888,899 | 562 | 4:5,063,973-5,064,343 | 374 | PCR, AFLP | This study |
| Marker 2d | GTTGCCAAAAGAGAGTTGACTG | TGACAAAAAGTATTAAAGGCAAGC | III:4,980,138-4,980,599 | 462 | 4:5,172,017-5,172,359 | 346 | PCR, AFLP | This study |
| Marker 2f | TACTTTTGTCTCCACACTTCATGG | CCCCTCTAGATTGTACATTGAAAGC | III:5,031,318-5,031,600 | 283 | 4:5,214,289-5,214,522 | 237 | PCR, AFLP | This study |
| Marker 2i | ATATTGAAAACTGTGATGGGGTTG | TGTCGATTTCTGAAAGTGCTACAG | III:5,515,111-5,515,400 | 290 | 4:5,709,055-5,709,270 | 223 | PCR, AFLP | This study |
| Marker 4a | GGGAATTAAAAGTTTCGGAGATTC | CCTGGCTTCTGATCACTGTTACAC | III:6,735,708-6,736,005 | 298 | 4:6,788,115-6,788,310 | 197 | PCR, AFLP | This study |

**Table S6**. Sequencing metrics for runs used in this study.

| Sequencing run | Sequencing strategy | Sequencing method | Sequencing platform | Extraction method | Library prep | Used for |
|---|---|---|---|---|---|---|
| HK104_longread_WGS2 | Whole-genome sequencing of *C. briggsae* wild isolate HK104 | Oxford Nanopore | PromethION | Phenol/chloroform | Ligation Sequencing Kit (ONT) | Building HK104 genome |
| QR24_longread_WGS | Whole-genome sequencing of *C. briggsae* wild isolate QR24 | Oxford Nanopore | PromethION | Phenol/chloroform | Ligation Sequencing Kit, Native Barcoding Kit (ONT) | Locating copies of *msft-1/tlpr-1* TA element |
| ED3036_longread_WGS | Whole-genome sequencing of *C. briggsae* wild isolate ED3036 | Oxford Nanopore | PromethION | Phenol/chloroform | Ligation Sequencing Kit, Native Barcoding Kit (ONT) | Locating copies of *msft-1/tlpr-1* TA element |
| JU439_longread_WGS | Whole-genome sequencing of *C. briggsae* wild isolate JU439 | Oxford Nanopore | PromethION | Phenol/chloroform | Ligation Sequencing Kit, Native Barcoding Kit (ONT) | Locating copies of *msft-1/tlpr-1* TA element |
| SB355_longread_WGS | Whole-genome sequencing of *C. plicata* SB355 | Oxford Nanopore | PromethION | Phenol/chloroform | Ligation Sequencing Kit (ONT) | Building *C. plicata* genome |
| HK104->AF16_16x MaternalBC_WGS | Whole-genome sequencing of *C. briggsae* HK104->AF16 maternal backcross (16x) | Illumina | MiSeq PE75 | Lucigen MasterPure | Nextera DNA Flex kit | Mapping HK104 introgression |
| INK199_WGS | Whole-genome sequencing of *C. briggsae* HK104->AF16 NIL1 | Illumina | NextSeq550 PE75 Medium | Lucigen MasterPure | Nextera DNA Flex kit | Mapping coordinates of Chr. III introgression |
| INK200_WGS | Whole-genome sequencing of *C. briggsae* HK104->AF16 NIL2 | Illumina | NextSeq550 PE75 Medium | Lucigen MasterPure | Nextera DNA Flex kit | Mapping coordinates of Chr. III introgression |
| PB1113_WGS | Whole-genome sequencing of *C. briggsae* AF16/HK104 AI-RIL | Illumina | MiSeq PE75 | Lucigen MasterPure | Nextera DNA Flex kit | Narrowing candidate region |
| PB1151_WGS | Whole-genome sequencing of *C. briggsae* AF16/HK104 AI-RIL | Illumina | MiSeq PE75 | Lucigen MasterPure | Nextera DNA Flex kit | Narrowing candidate region |
| PB1171_WGS | Whole-genome sequencing of *C. briggsae* AF16/HK104 AI-RIL | Illumina | NextSeq550 PE75 Medium | Lucigen MasterPure | Nextera DNA Flex kit | Narrowing candidate region |
| AF16_RNAseq | Transcriptome sequencing of C. briggsae reference AF16 on mixed-stage embryos | Illumina | NextSeq550 PE75 Medium | TRIzol/chloroform | TruSeq RNA Library Prep kit | Identifying candidate genes |
| HK104_RNAseq | Transcriptome sequencing of C. briggsae wild isolate HK104 on mixed-stage embryos | Illumina | NextSeq550 PE75 Medium | TRIzol/chloroform | TruSeq RNA Library Prep kit | Identifying candidate genes, annotating HK104 genome |
| SB355_WGS | Whole-genome sequencing of C. plicata SB355 | Illumina | NextSeq550 PE150 Medium | Lucigen MasterPure | NEBNext Ultra library prep kit | Polishing *C. plicata* genome |
| SB355_RNAseq | Transcriptome sequencing of C. plicata reference SB355 on L4-stage larvae | Illumina | NextSeq2000 P2 SR100 | TRIzol/chloroform; NEBNext rRNA Depletion kit | NEBNext Multiplex Unique Dual Indexes | Annotation of *C. plicata* genome |
